# Dutch population structure across space, time and GWAS design

**DOI:** 10.1101/2020.01.01.892513

**Authors:** Ross P Byrne, Wouter van Rheenen, Project MinE ALS GWAS Consortium, Leonard H van den Berg, Jan H Veldink, Russell L McLaughlin

**Author notes:** A list of Project MinE ALS GWAS Consortium authors and affiliations appears in Supplementary note 1. To whom correspondence should be addressed: Ross P Byrne and Russell L McLaughlin, Complex Trait Genomics Laboratory, Smurfit Institute of Genetics, Trinity College Dublin, Dublin D02 DK07, Republic of Ireland.

## Abstract

We studied fine-grained population genetic structure and demographic change across the Netherlands using genome-wide single nucleotide polymorphism data (1,626 individuals) with associated geography (1,422 individuals). We applied ChromoPainter/fineSTRUCTURE, identifying 40 haplotypic clusters exhibiting strong north/south variation and fine-scale differentiation within provinces. Clustering is tied to country-wide ancestry gradients from neighbouring lands and to locally restricted gene flow across major Dutch rivers. Despite superexponential population growth, north-south structure is temporally stable, with west-east differentiation more transient, potentially influenced by migrations during the middle ages. Within Dutch and international data, GWAS incorporating fine-grained haplotypic covariates are less confounded than standard methods.

---

The Netherlands is a densely populated country on the northwestern edge of the European continent, bounded by Germany, Belgium and the North Sea. The country is divided into twelve provinces and has a complex demographic history, with occupation by several Germanic peoples since the collapse of the Roman Empire, including the Frisians, the Low Saxons and the Franks. Over 17 million individuals now inhabit this relatively small region (41,500km^2^), making it one of the most densely populated countries in Europe. Despite its small geographical size, previous genetic studies of the people of the Netherlands have demonstrated coarse population structure that correlates with its geography, as well as apparent heterogeneity in effective population sizes across provinces^1,2^. These observations suggest that the demographic past of the Dutch population has left residual signatures in its present regional genetic structure; however this has not been fully explained in the context of neighbouring populations and thus far the use of unlinked genetic markers have limited the resolution at which this structure can be described. This resolution limit also confines the extent to which the confounding effects of population structure can be controlled in genomic studies of health and disease such as genome-wide association studies (GWAS). As these studies continue to seek ever-rarer genetic variation with ever-increasing cohort sizes, intricate understanding and fine control of population structure is becoming increasingly relevant, but increasingly challenging^3^.

Recent studies have showcased the power of leveraging shared haplotypes to uncover and characterise previously unrecognised fine-grained genetic structure within populations, yielding novel insights into the demographic composition and history of Britain and Ireland^4–7^, Finland^8^, Japan^9^, Italy^10^ and Spain^11^. Haplotype sharing has also revealed genetic affinities between populations, enabling inference of historical admixture events using modern populations as proxies for ancestral admixing sources^12^. Furthermore, geographic information can be integrated to model genetic similarity as a function of spatial distance^13^ to infer demographic mobility within or between populations; one approach uses the Wishart distribution to estimate and map a surface of effective migration rates based on deviations from a pure isolation by distance model^14^, allowing migrational cold spots to be inferred which may derive from geographical boundaries such as rivers and mountains. Almost half of the area of the Netherlands is reclaimed from the sea and its contemporary land surface is densely subdivided by human-made waterways and naturally-occurring rivers, including the Rhine (Dutch: *Rijn*), Meuse (*Maas*), Waal and IJssel. These rivers have been speculatively linked to genetic differentiation between northern and southern Dutch subpopulations in previous work^1^; however the explicit relationship between Dutch genetic diversity and movement of people within the Netherlands has not been directly modelled.

The Dutch have previously received special interest as a model population^1,2^ and form a major component of substantial ongoing efforts to better understand human health, disease, demography and evolution. For example, at the time of writing, over 10% of all studies listed in the NHGRI-EBI genome-wide association study (GWAS) catalogue^15^ include the Netherlands in their “Country of recruitment” metadata. As well as offering insights into demography and human history, refined population genetic studies are important to identify and adequately control confounding effects in genomic studies of health and disease, especially if rare variants or spatially structured environmental factors contribute substantially to variance in phenotype^16^. In this study, we harness shared haplotypes to examine the fine-grained genetic structure of the Netherlands. We show that Dutch population structure is much stronger than previously recognised, and is ancient and persistent over time. The strength and stability of the observed structure appears to be tied to the relationship of the Netherlands to neighbouring lands and to its own internal geography, and has likely been shaped over history by migration, but preserved in recent generations by enduring sedentism of genetically similar individuals within regions. This has led to genetic structure that demonstrably confounds GWAS; however through analysis of the Netherlands and more extensive international data^17^, we show that using shared haplotypes as GWAS covariates significantly reduces this confounding over standard single-marker methods.

## Results

### The genetic structure of the Dutch population

We mapped the haplotypic coancestry profiles of 1,626 Dutch individuals using ChromoPainter^18^ and clustered the resulting matrix using fineSTRUCTURE^18^, identifying 40 genetic clusters at the highest level of the hierarchical tree which segregated with geographical provenance. We explored the clustering from the finest (k=40) to the coarsest level (k=2), settling on k=16 as it captured the major regional splits sufficiently with little redundancy. Clusters at this level were robustly defined by total variation distance (TVD) and fixation index (F_ST_; Figure 1a); remarkably, some F_ST_ values between Dutch clusters were comparable in magnitude to estimates between European countries (calculated using data from reference 19; Supplementary table 1). Some clusters had expansive geographical ranges (for example NHFG, representing individuals from North Holland, Friesland and Groningen), while others neatly distinguished populations on a sub-provincial level (for example, NBE and NBW, representing east and west regions of North Brabant). For visualisation we projected the ChromoPainter coancestry matrix in lower dimensional space using principal component analysis (PCA; Figure 1b) and assigned cluster labels based on majority sampling location (available for 1,422 individuals), arranging neighbouring and genetically similar clusters into cluster groups, as with previous work^6^. The first principal component (PC) of coancestry followed a strong north-south trend (latitude vs mean PC1 per town r^2^ = 0.52; p = 6.8×10^−72^) with PC2 generally explained by a west-east gradient (longitude vs mean PC2 per town r^2^ = 0.29; p = 3.4×10^−33^).

**Figure 1.**
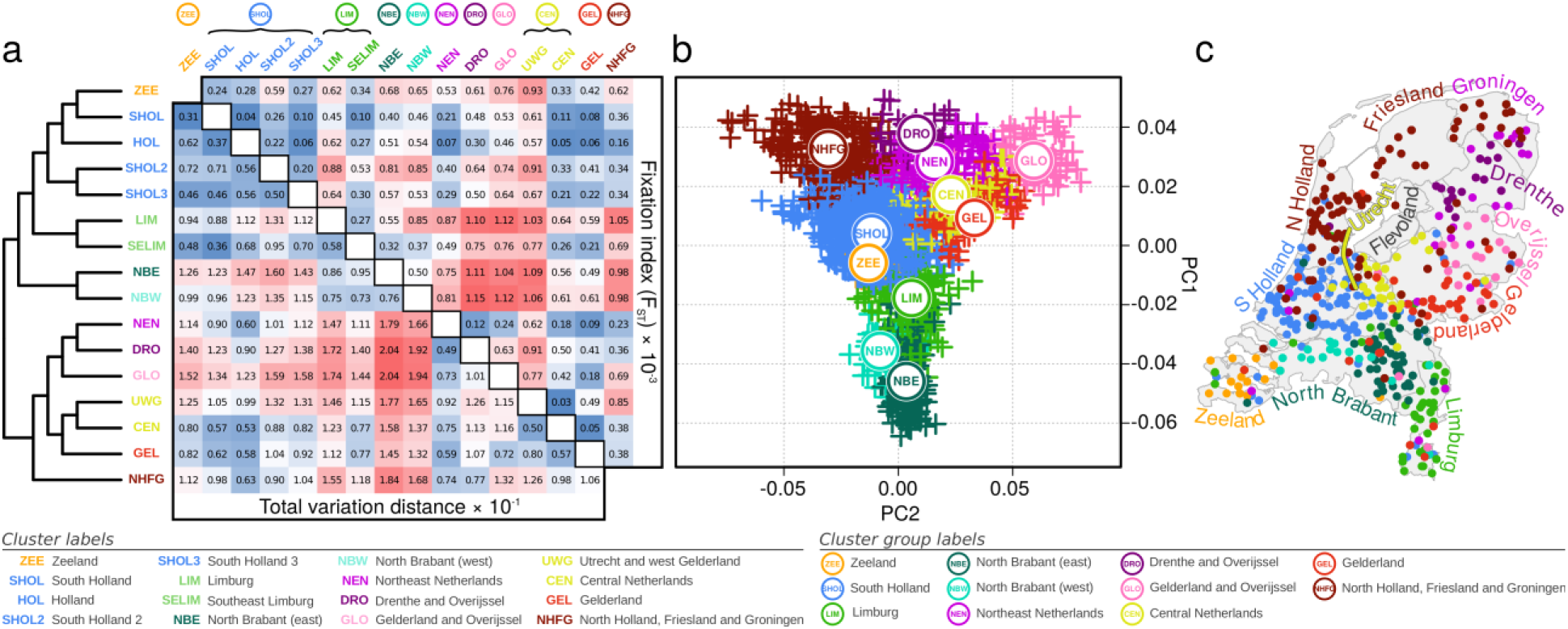
The genetic structure of the people of the Netherlands. (**a**) fineSTRUCTURE dendrogram of ChromoPainter coancestry matrix showing clustering of 1,626 Dutch individuals based on haplotypic similarity. Associated total variation distance (TVD) and fixation index statistics between clusters are shown in the matrix. Permutation testing of TVD yields p<0.001 for all cluster pairs, indicating that clustering is non-random. Cluster labels derive from Dutch provinces and are arranged into cluster groups for genetically and geographically similar clusters (circled labels). (**b**) The first two principal components (PCs) of ChromoPainter coancestry matrix for all individuals analysed. Points represent individuals and are coloured and labelled by cluster group. (**c**) Geographical distribution of 1,422 sampled individuals, coloured by cluster groups defined in (a). Labels represent provinces of the Netherlands. Map boundary data from the Database of Global Administrative Areas (GADM; https://gadm.org).

As previously observed in different populations^6^, the distribution of individuals in this genetic projection generally resembled their geographic distribution (Figure 1c), with some exceptions. For example, North Brabant is geographically further north than Limburg, but is further separated by PC1 from northern clusters. We explored the possibility that this could instead be explained by relative ancestral affinities to neighbouring lands by modelling the genome of each Dutch individual as a linear mixture of European sources (obtained from reference 19) using ChromoPainter, retaining source groups that best matched Dutch individuals for at least 5% of the genome^4^ (Figure 2). The resulting profiles of German, Belgian and Danish ancestries were significantly autocorrelated (p_DE_, p_BE_ < 0.0001; p_DK_ < 0.001; Moran’s I and Mantel’s test) and spatially arranged along geographical directions S66°W, N73° E and S73°E respectively, approximately corresponding to declining ancestry gradients directed away from the German and Belgian borders and the North Sea boundary (Figure 2; 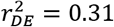; 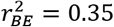; 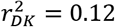; *p*_*DE*_ = 9.4 ×10^−119^; *p*_*BE*_ = 2.7 ×10^−133^; *p*_*DK*_ = 1.1 × 10^−39^). The spatial distribution of French ancestry was comparatively uniform, with only a modest correlation due east (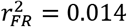; *p*_*FR*_ = 9.5 × 10^−6^). The general trend across the Netherlands was thus of complementary Belgian and German ancestral affinities, decaying with distance from the respective borders. North Brabant, however, showed a greater Belgian profile than Limburg, despite similar, substantial Belgian frontiers in both Dutch provinces. Conversely, the German ancestry profile of Limburg greatly exceeded that of North Brabant, reflecting its 200-kilometre border with Germany and centuries of consequent demographic contact and likely genetic admixture.

**Figure 2.**
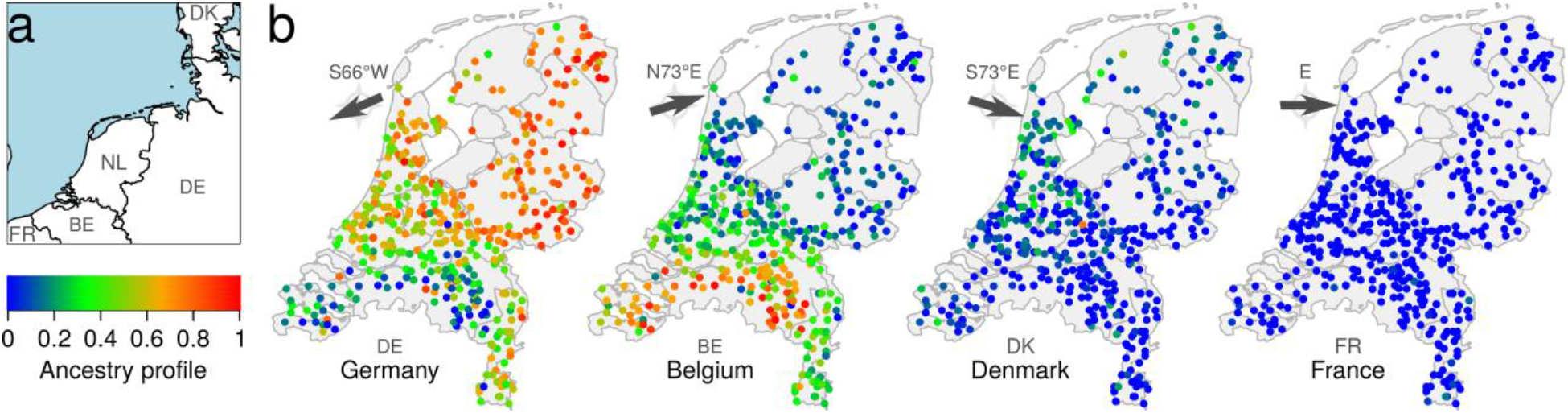
The ancestry profile of the Netherlands. (**a**) The Netherlands and its geographical relationship to neighbouring lands. (**b**) German, Belgian, Danish and French haplotypic ancestry profiles for 1,422 Dutch individuals. Arrows indicate the predominant directions along which the ancestry gradients are arranged across the Netherlands. Map boundary data from the Database of Global Administrative Areas (GADM; https://gadm.org) and Natural Earth (https://naturalearthdata.com).

### Genome flux and stasis in the Netherlands

To explore temporal trends in Dutch population structure we called genomic segments of pairwise identity-by-descent (IBD) using RefinedIBD^20^. An IBD haplotype sharing matrix is conceptually similar to a ChromoPainter coancestry matrix^21^, but trades some sensitivity to be more explicitly interpretable. As IBD segment length is inversely related to age^22,23^, different length intervals can inform on structure at different time depths. Total pairwise IBD between Dutch individuals mirrored the structure observed with ChromoPainter (Figure 3a), with 8 distinct clusters identified in the IBD sharing matrix that broadly segregated with geography and recapitulated some of the important splits obtained from fineSTRUCTURE, most strikingly the west-east split in North Brabant. Decomposing total IBD by centiMorgan (cM) length into short (1-3 cM), medium (3-5 cM) and long (5-7 cM) bins, we observed stability over time of north-south structure and the emergence of west-east structure embedded in 3-5 cM segments (Figure 3b), corresponding to an expected time depth around 1,120 years ago^23^. As this date and the structure observed is dependent on the (arbitrary) thresholds set for IBD segment length bins, we have also provided an interactive environment in which Dutch population structure can be explored across a range of IBD segment bins (bioinf.gen.tcd.ie/ctg/nlibd).

**Figure 3.**
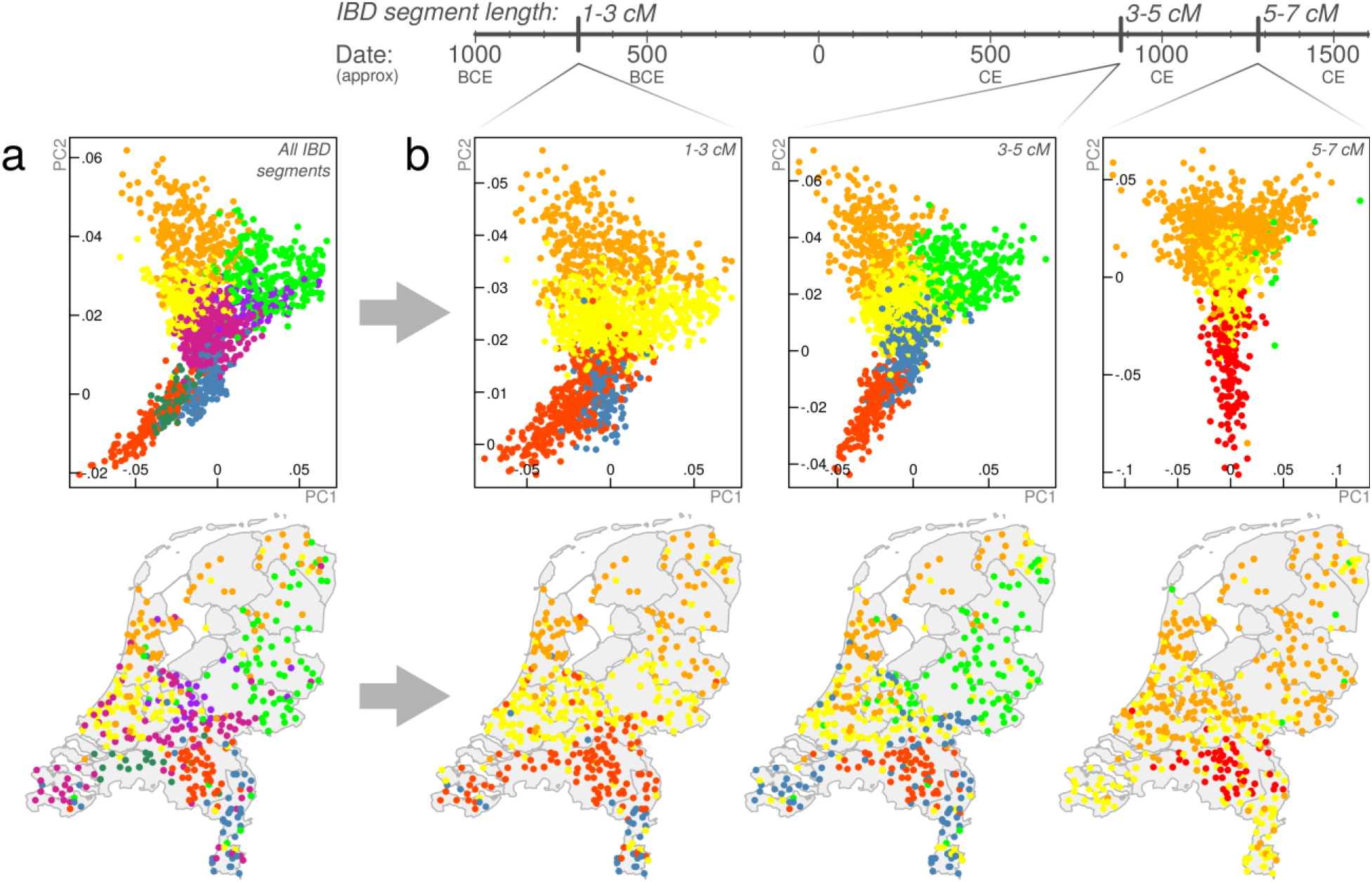
The changing genomic structure of the Dutch population over time. (**a**) Principal component (PC) analysis of pairwise total identity-by-descent (IBD) for 1,626 Dutch individuals (top) and their geographical provenance (bottom). Points represent individuals and are coloured by cluster assignment (mclust on pairwise IBD matrix). (**b**) PCs (top) and geographical provenance (bottom) for pairwise sharing of 1-3, 3-5 and 5-7 centiMorgan (cM) IBD segments, corresponding to point estimates of expected time depths at approximately 2,700, 1,120 and 720 years ago, respectively. Time depths for IBD segment bins have wide distributions^23^; expected values presented here should be interpreted as a guide only and the changing west-east structure over time does not necessarily reflect (for instance) a precisely-timed admixture event. Map boundary data from the Database of Global Administrative Areas (GADM; https://gadm.org).

Although these observations could potentially be biased by power to detect population structure in longer and shorter bins, the temporally volatile west-east structure contrasts with the stability and persistence of old north-south structure and possibly represents a genomic signature of historical demographic flux in the region and its surrounding lands. With this in mind, we investigated possible admixture from outside demographic groups using GLOBETROTTER^12^ with 4,514 European individuals^19^ representing modern proxies for admixing sources. Across the Dutch sample, significant migration dating to 1088 CE (95% c.i. 1004-1111 CE) was inferred with the major contributing source best modelled by modern Germans and the minor source best modelled by southern European groups (France, Spain) (Table 1). This is supported by single-marker ADMIXTURE component estimates showing that the Netherlands has the closest profile to Germanic groups (Supplementary figure 1) and is consistent with the ancestry profile gradients detailed in Figure 2. The timing of the inferred 11^th^ century event was stable across Dutch fineSTRUCTURE clusters (to varying degrees of confidence), suggesting that the signal represents an important period in the establishment of the modern Dutch genome (Table 1); however, given the state of demographic flux in Europe at the time, its exact historical correlate is open to interpretation. Notably, a significant admixture event with a major Danish source was inferred between 759 and 1290 CE in the NHFG cluster group (representing Dutch northern seaboard provinces); this period spans a historical period of recorded Danish Viking contact and rule in northern Dutch territories.

**Table 1.**
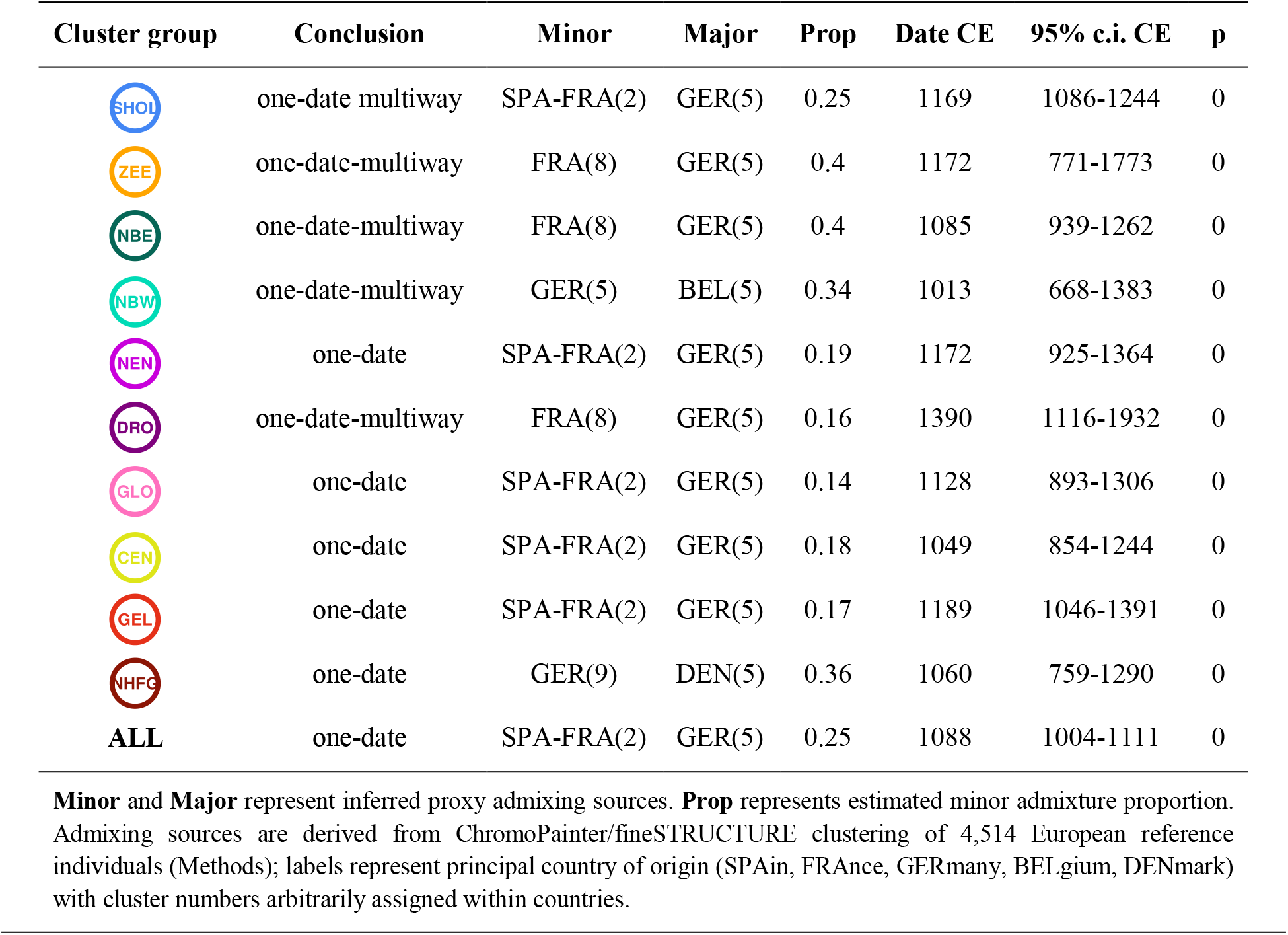
GLOBETROTTER date and source estimates for admixture into the Netherlands.

In addition to influence from outside populations, the population structure detailed in Figure 1 and Figure 3 has likely been shaped by independent demographic histories within the Netherlands. In support of this, we noted that short (1-2 cM) IBD segments shared between northern clusters and provinces outnumbered those shared between southern clusters and provinces (Supplementary figure 2), and, as observed previously^2^, northern provinces shared more short segments with southern provinces than southern provinces shared amongst themselves. Together, these results suggest that the north had a smaller ancestral effective population size (N_e_) than the south and is probably derived from an ancient or historical founder event forming the northern population from a subset of southerners. We formally characterised ancestral trajectories in N_e_ between the north and the south of the Netherlands using the nonparametric method IBDNe^24^ for the entire Dutch sample and two subsamples representing the principal fineSTRUCTURE north/south split (Figure 4a), retaining a random sample of 641 individuals from each group. We also characterised historical N_e_ within individual Dutch provinces for which genotypes for more than 40 individuals were available. Countrywide, N_e_ has grown superexponentially over the past 50 generations in the Netherlands (Figure 4a) and has been consistently lower in the north than the south. Despite this, the pattern of growth in northern and southern groups was identical, with a steady exponential growth up to around 1650 CE, when a major uptick in growth rate was observed. This corresponds to a period of substantial economic development in the Netherlands over the 17^th^ century known to historians as the Dutch Golden Age. Preceding this period, historical N_e_ estimates for the entire country and for northern/southern groups showed only a modest response to the Black Death (*Yersinia pestis* plague pandemic) of the 14^th^ century which claimed up to 60% of Europe’s population^25^. Conversely, N_e_ estimation within individual Dutch provinces revealed a much more detectable impact (Figure 4b).

**Figure 4.**
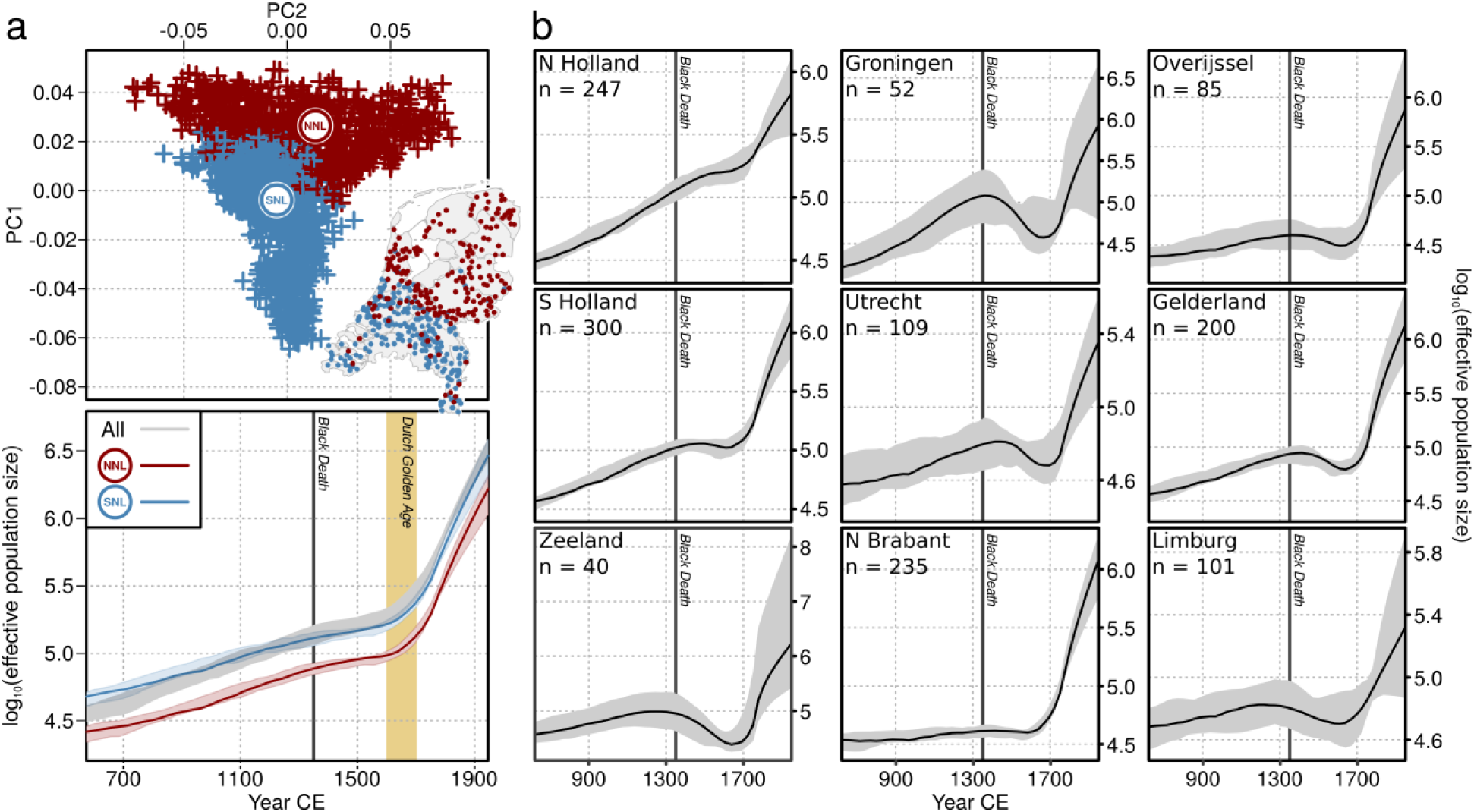
Dutch effective population size over time. (**a**) Historical change in effective population size (N_e_) over the past 50 generations for all Dutch individuals and subsets of northerners and southerners. The top plot shows the principal components of ChromoPainter coancestry coloured by the first (k=2) fineSTRUCTURE split, which separates the Dutch population into northern (NNL) and southern (SNL) genetic clusters; inset shows geographical distribution of these individuals. The bottom plot shows growth in effective population size countrywide or per fineSTRUCTURE cluster over the past 50 generations. (**b**) Historical N_e_ trajectories for individual Dutch provinces with more than 40 individuals sampled. N_e_ plots show estimates ± 95% c.i. and assume 28 years per generation and mean year of birth at 1946 CE. Map boundary data from the Database of Global Administrative Areas (GADM; https://gadm.org).

### Genomic signatures of Dutch mobility

We noted that long (>7 cM) IBD segments, which capture recent shared ancestry, were almost always shared within genetic clusters (and provinces), and rarely between (Supplementary figure 2). This indicates a propensity for genetically similar individuals (relatives) to remain mutually geographically proximal, suggesting a degree of sedentism that has likely influenced Dutch population structure over time. It has also previously been argued that genetic structure in the Netherlands may be partially rooted in geographic obstacles imposed by the country’s major waterways^1^ so we explicitly modelled genetic similarity as a function of geographic distance using EEMS^14^ to infer migrational hot and cold spots (Figure 5). The resulting effective migration surface showed several apparent barriers to gene flow, the strongest and most contiguous of which runs in an east-west direction across the Netherlands overlapping the courses of the Rhine, Meuse and Waal rivers. This inferred migrational boundary also approximately corresponds to the geographical division determining the principal fineSTRUCTURE split between northern and southern Dutch populations (Figure 4a) as well as the geographical boundaries between clusters inferred from ancient IBD segments (Figure 3b), suggesting that these rivers have been a historically persistent determinant of Dutch population structure.

**Figure 5.**
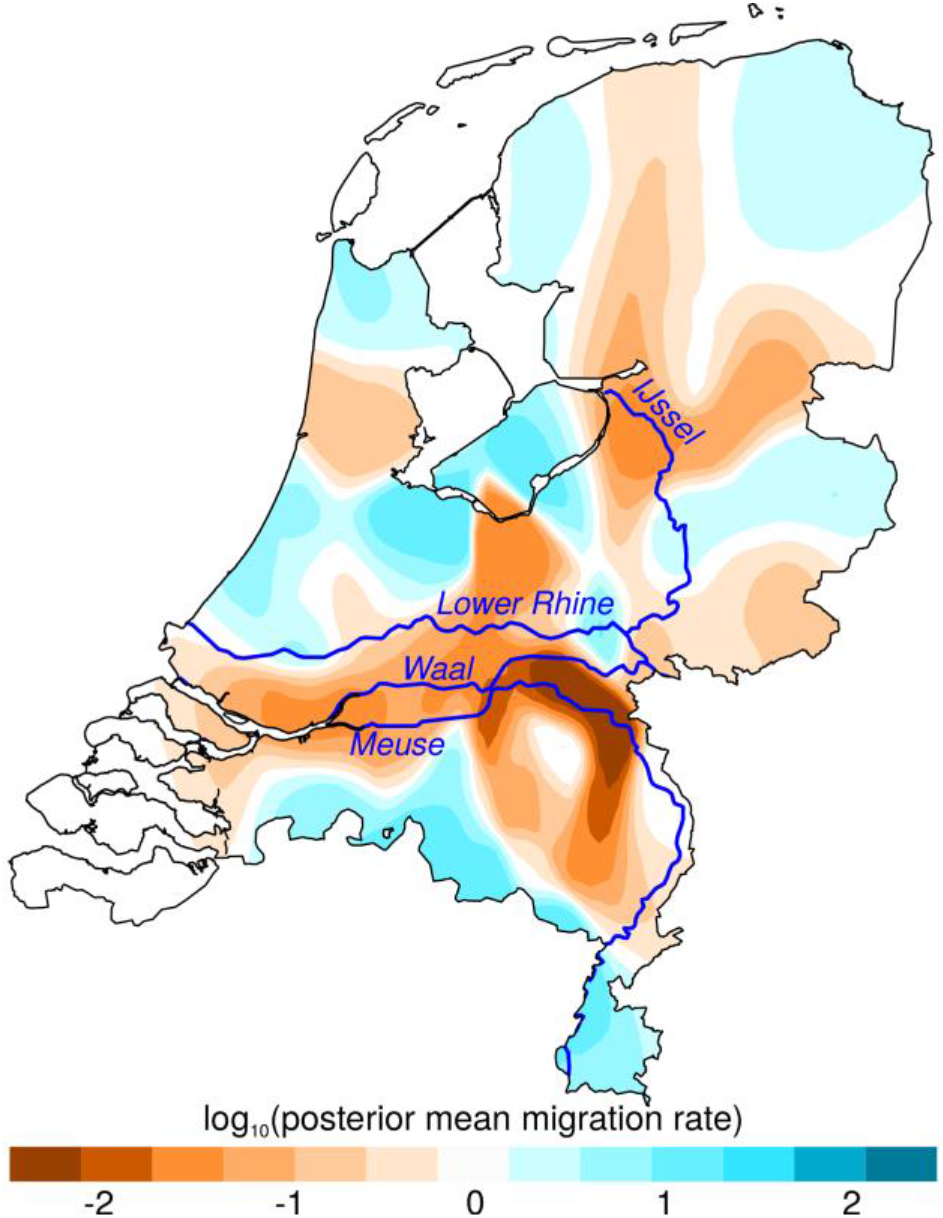
The effective migration surface of the Netherlands. Contour map shows the mean of 10 independent EEMS posterior migration rate estimates between 800 demes modelled over the land surface of the Netherlands. A value of 1 (blue) indicates a tenfold greater migration rate over the average; −1 (orange) indicates tenfold lower migration than average. The courses of major rivers are included to highlight their correlation with migrational cold spots. Map boundary data from the Database of Global Administrative Areas (GADM; https://gadm.org); river course data from Natural Earth (https://www.naturalearthdata.com).

### GWAS confounding by fine-grained structure

As population structure confounds GWAS (for example due to stratification of cases and controls between subpopulations), we investigated the extent to which haplotype sharing captures confounding structure in a Dutch sample of 1,971 cases of amyotrophic lateral sclerosis (ALS) and 2,782 controls from a recent multi-population GWAS for ALS^17^. PCs of the haplotypic ChromoPainter coancestry matrix for these 4,753 individuals explained substantially more variance in ALS phenotype than PCs calculated from SNP genotypes alone, indicating latent structure captured by ChromoPainter that is stratified between cases and controls (Figure 6a). To estimate the extent to which this stratified structure confounds GWAS we calculated case-control association statistics using a logistic model covarying for either 20 ChromoPainter PCs or 20 SNP PCs and estimated the linkage disequilibrium (LD) score regression intercepts for both sets of resulting summary statistics. An intercept higher than 1 indicates confounding in the GWAS; Figure 6a shows that GWAS statistics calculated with ChromoPainter PCs as covariates are less confounded than statistics using SNP PCs, albeit with overlapping confidence intervals for the relatively small Dutch sample. To more adequately represent the large-scale multi-population data typically used in modern GWAS, we extended our analysis to the full ALS case-control dataset from which the Dutch data derive^17^, including 36,052 individuals from twelve European countries and the USA. For computational tractability, instead of ChromoPainter we used PBWT-paint (https://github.com/richarddurbin/pbwt/blob/master/pbwtPaint.c), a scalable approximate haplotype painting method based on the positional Burrows-Wheeler transform^26^. When run on our original Dutch dataset of 1,626 individuals, the structure rendered by PBWT-paint was almost identical to ChromoPainter (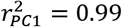; 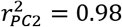; Supplementary figure 3), indicating its suitability for this analysis. PBWT-paint captured pervasive global and local structure in the multi-population GWAS data that both separated and subdivided countries (Figure 6b). Top PCs of PBWT-paint coancestry explained substantially more variance in phenotype than SNP PCs and GWAS statistics including PBWT-paint PCs as covariates were significantly less confounded than statistics corrected by SNP PCA (Figure 6a, LD score regression intercepts).

**Figure 6.**
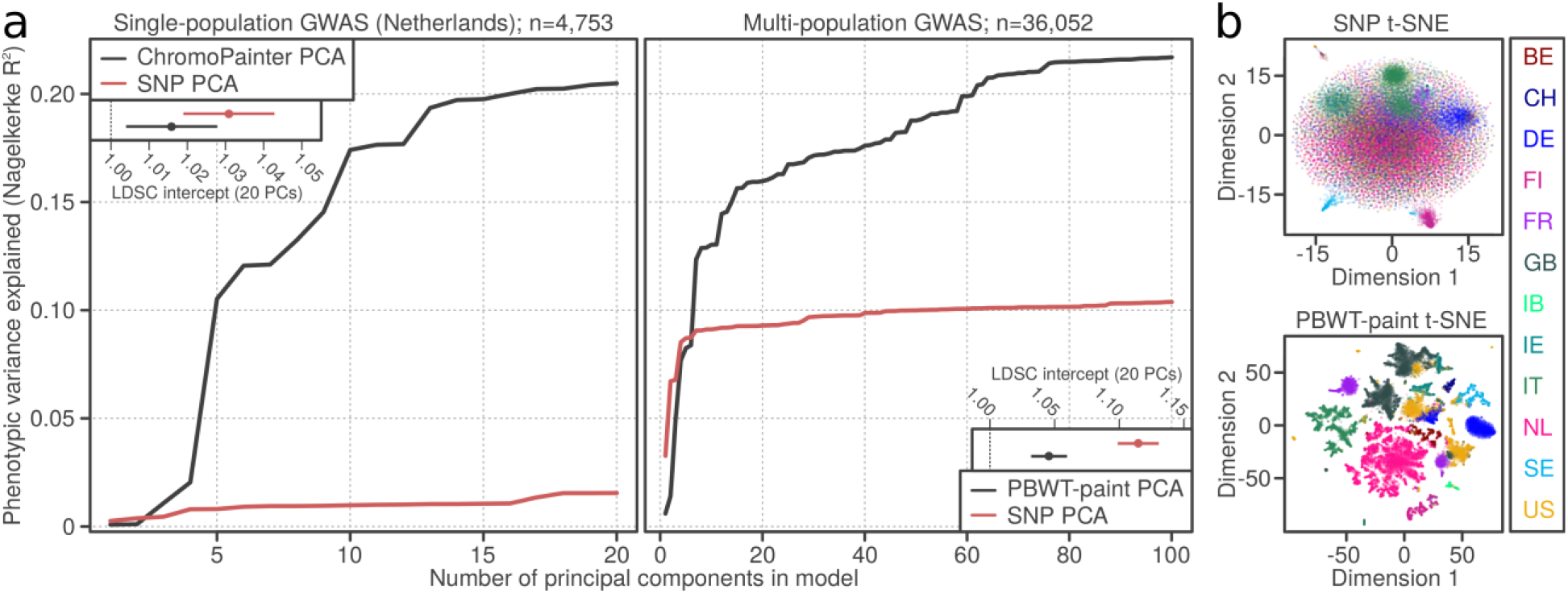
Fine-grained population structure and genome-wide association study (GWAS) confounding. (**a**) Variance in phenotype (amyotrophic lateral sclerosis) explained by principal components (PCs) for a single-population Dutch GWAS (left) and a multi-population GWAS (right). Insets show linkage disequilibrium score regression (LDSC) intercept terms (a summary estimate of GWAS confounding) when the first 20 single nucleotide polymorphism (SNP)-based PCs (SNP PCA) or the first 20 haplotype-based PCs (ChromoPainter/PBWT-paint PCA) are included as GWAS covariates. (**b**) Summary visualisations (t-distributed stochastic neighbour embedding, t-SNE) of local and global structure in the multi-population GWAS based on SNP genotypes (top) or haplotype sharing inferred using the scalable PBWT-paint chromosome painting algorithm (bottom). Individuals are coloured by country of origin; labels (right) follow ISO 3166-1 country codes, except IB, which was labelled Iberia (containing Spanish and Portuguese data) in the original GWAS dataset. PCA, principal component analysis; PBWT, positional Burrows-Wheeler transform.

## Discussion

We have studied the Netherlands as a model population, harnessing information from shared haplotypes and recent developments in spatial modelling to gain intricate insights into the geospatial distribution and likely origin of Dutch population genetic structure. The structure identified through shared haplotypes is surprisingly strong; some Dutch genetic clusters identified this way are more mutually distinct (by F_ST_) than whole European countries. We have also introduced a novel use of IBD sharing combined with PCA and Gaussian mixture model-based clustering to characterise changing population structure over time, revealing transient genetic structure layered over strong and stable north-south differentiation in the Netherlands. This is contextualised by somewhat distinct demographic histories between genetic groups in the Netherlands, with consistently lower N_e_ in the north than the south. A potential source of the north-south differentiation is impaired migration across the east-west courses of the Rhine, Meuse and Waal, which effectively separate southern Dutch populations from the north. The population structure observed in the Netherlands is especially remarkable when considered in terms of the country’s size and extensive infrastructure; notably Denmark, which is roughly equal in geographical area, is genetically homogeneous, forming only a single cluster when interrogated using fineSTRUCTURE^27^, despite its island-rich geography. Both the United Kingdom and Ireland also exhibit at least one large indivisible cluster constituting a large fraction of the population^4–6^, however no extraordinarily large clusters dominate the Dutch sample. Mean F_ST_ between Dutch clusters also greatly outmeasures that observed between Irish clusters, suggesting that the extent of population differentiation is higher in the Netherlands, despite Dutch land area being less than half that of the island of Ireland.

While coarse geographical trends in Dutch genetic structure have previously been described using single-marker PCA^1^, our use of shared haplotypes reveals structure at a much higher resolution, differentiating subpopulations between, and sometimes within, provinces (Figure 1). As a striking example, individuals from the east and west of North Brabant (NBE and NBW in Figure 1) are mutually genetically distinguishable and are more distinct from clusters to their north than Limburg, despite being geographically closer. This deviation from haplotype sharing mirroring geography appears to be driven by strong genetic affinity to Belgium (Figure 2), reflecting a long history of demographic and sovereign overlap across a 100 km frontier spanning the modern Dutch-Belgian border. In contrast, the majority of ancestral influence in Limburg, which also shares a substantial border with Belgium, is equally split between Belgium to the west and Germany to the east. Notably, the Belgian border with the south of Dutch Limburg is almost entirely described by the course of the Meuse, which may have acted as a historical impediment to migration, thus distinguishing individuals in this region genetically. This is reflected in IBD clustering, in particular the distinction of southern Limburgish individuals from the rest of the Netherlands in short (1-3 cM) segments, which otherwise only describe coarse north-central-south structure (Figure 3). Future work explicitly modelling Dutch-Belgian and Dutch-German frontiers using additional Belgian and German genetic data with associated geography will resolve the historical and present-day role of the Meuse in distinguishing distinct population clusters in the south of the Netherlands.

Similarly to North Brabant, groups of individuals in North and South Holland show significant genetic separation despite mutual geographic proximity. While we have chosen to group the four South Holland clusters for visual brevity in Figure 1, they are robustly distinct by TVD analysis (Figure 1a), indicating that significant population differentiation exists even within South Holland. Migration and admixture in the highly urbanised *Randstad* has been proposed as a driver of genetic diversity and loss of geographic structure in this region^1^; the overlaid geographical distribution of regional ancestry profiles (Figure 2) for this area lends support to this hypothesis. However, the geographical ranges of the four South Holland clusters are somewhat independent (Supplementary figure 4), indicating that some degree of genetic structure has survived this urbanisation. Previous studies have highlighted the correlation between decreasing autozygosity and increased urbanisation^28^; future work leveraging the ChromoPainter/fineSTRUCTURE framework coupled with length-binned IBD and Gaussian mixture model-based clustering will more explicitly delineate the interplay between urbanisation and population structure over time. To this end, highly urbanised areas such as the *Randstad* will be particularly informative.

The principal fineSTRUCTURE split in the Netherlands describes north-south genetic differentiation (Figure 1) that is strong and persistent over time (Figure 3). We hypothesised that this reflects partially independent demographic histories so we estimated ancestral N_e_ for northern (NNL) and southern (SNL) Dutch fineSTRUCTURE populations, revealing superexponential growth in both populations with a sudden increase in rate following the 17^th^ century Dutch Golden Age (Figure 4a). Historical N_e_ follows the same approximate trajectory for both populations but is consistently lower for the northern cluster, corroborating previous observations of increased homozygosity in northern Dutch populations^1^ and consistent with a model of northerners representing a founder isolate from southerners (although a more complex demographic model may better explain these observations)^1,2^. The apparent absence of N_e_ decline in 14^th^-century Netherlands initially hints at the possibility that the Black Death had a weaker impact in the region than elsewhere in Europe; although this agrees with the views of some historians, it is hotly debated by others^29^. Per province, however, most N_e_ estimates display a prominent dip at this time (Figure 4b), suggesting that merging non-randomly mating subpopulations into a countrywide group (Figure 4a) artificially inflates diversity, thus smoothing over any population crash following the Black Death. Population structure is thus important when estimating N_e_ and trends countrywide and in NNL and SNL clusters (Figure 4a) should be interpreted carefully: it is possible that a substantial population crash brought about by the Black Death might have had only a marginal impact on the overall effective size of the breeding population in these merged groups. Indeed, the rate of exponential growth in countrywide N_e_ (Figure 4a) is marginally shallower in the 10 generations following the Black Death (0.024; 95% c.i. 0.0235-0.0251) compared to the 10 generations prior (0.017; 95% c.i. 0.016-0.018), indicating enduring strain on the overall Dutch population prior to its recovery in the Dutch Golden Age.

Previous works have hinted that north-south genetic differentiation in the Netherlands may have been facilitated by cultural division between the predominantly Catholic south and the Protestant north^1^. Given that the north-south structure observed in 1-3 cM IBD bins (expected time depth ~700 BCE) greatly precedes different forms of Christianity (Figure 3), our data support a model in which the Protestant Reformation of the 16^th^ and 17^th^ centuries exploited pre-existing demographic subdivisions, leading to correlation between distinct cultural affinities and clusters of genetic similarity. Geographical modelling supports the role of migrational boundaries in establishing and maintaining this population substructure, especially rivers (Figure 5). A substantial belt of low inferred migration runs across the Netherlands, corresponding closely to the roughly parallel east-west courses of the Lower Rhine, Waal and Meuse rivers and correlating with the geographical boundary of the principal north-south fineSTRUCTURE split. Absolute assignment of causality to these geographical correlates is, however, not possible and, given the dense network of waterways in the Netherlands, could be misleading. For example, a strong migrational cold spot in the east of the Netherlands runs parallel to the IJssel (Figure 5), but could potentially be better explained by the course of the Apeldoorn Canal, a politically fraught waterway constructed in the early 19^th^ Century. Similarly, a cold spot in the northwest directly overlays the North Sea Canal (completed in 1876). As both of these are human-made waterways, it is not certain whether their courses are consequences or determinants of low movement of people across their paths.

As well as internal geography, outside populations have also played an important and significant role in the establishment of population structure in the Netherlands (Figure 2; Table 1); however the variety and extent of demographic upheaval and mobility of European populations over history obscure the likely historical provenance of many inferred admixture signals. As an important exception, however, ancestry profiles show a small but significant contribution of Danish haplotypes in the north and west of the Netherlands, a possible vestige of Viking raids in coastal areas in the 9^th^ and 10^th^ centuries. This is corroborated by an inferred GLOBETROTTER single-date admixture event in the NHFG (North Holland, Friesland and Groningen) cluster (Figure 1) between 759 and 1290 CE with Danish haplotypes as a major admixing source (Table 1). The demographic legacy of more than a century of Danish Viking raids and settlement in the Netherlands has been the subject of some debate; from our data, it appears that the modern Dutch genome has indeed been partially shaped by historical Viking admixture. This Danish Viking contact is contemporaneous with a critical period in the establishment of the modern Dutch genome from other outside sources (1004-1111 CE; Table 1), although the precise historical correlates of the admixture events detected in the remaining Dutch regions are less obvious. Future densely sampled ancient DNA datasets from informative time depths in the Netherlands and northwest Europe will enable direct estimation of ancestral population structure, admixture, demographic affinities and effective population sizes, improving precision over the current study which depends on proxy patterns of haplotype sharing between modern individuals. Similarly, regional ancestry and admixture inference are limited by the use of modern proxy populations in place of true ancestral sources; nevertheless, there are ample advantages to the use of modern data, including large sample size and relevance to research on modern human health and disease. In particular, as in our previous work in Ireland^6^, samples in the current Dutch dataset were not specifically selected to have pure ancestry in each geographical area (eg all grandparents from the same region^4^) meaning the degree of structure observed is not idealised or exaggerated by sampling, but instead representative of the structure expected in any GWAS that includes Dutch data.

We therefore explored the impact of fine-scale genetic structure described in this study and others^4–11^ on GWAS statistics, using the ALS study from which the Dutch data derive as an exemplar trait. Generally, population-based PCs should not predict case/control status; if they do, this indicates that (sub)populations are stratified between cases and controls, introducing bias that artificially inflates GWAS statistics. In both Dutch-only and multi-population analyses, fine-scale genetic structure detected by haplotype sharing (ChromoPainter or PBWT-paint) explained substantially more variance in phenotype (ALS case/control status) than standard SNP-only PCA (Figure 6a). This demonstrates the power of shared haplotypes to simultaneously capture subtle genetic structure within single countries (that is potentially invisible to standard single-marker PCA), broader structure between countries and potential cryptic technical artefacts such as platform- or imputation-derived bias. We found that shared haplotypes are effective for controlling GWAS inflation: statistics calculated using haplotype-based PCs as covariates showed lower overall confounding than single marker-based covariates, as measured by LD score regression intercepts (Figure 6a). In the age of large-scale, single-country and cross-population biobanks, the additional power of haplotype sharing methods to detect fine-scale local population structure will be crucial for ensuring robust GWAS results unconfounded by ancestry. For example, a recent study of latent structure in the UK Biobank demonstrated that a GWAS for birth location returned significant hits even after correction for 40 single-marker PCs^30^, suggesting that residual fine-grained population structure may influence other GWAS from this cohort. Ongoing developments in scalable haplotype sharing algorithms such as PBWT-paint will help to address this problem by facilitating the creation of biobank-scale haplotype sharing resources, simultaneously improving studies of human health and disease and enabling large-scale, fine-grained population genetic studies of human demography.

## Methods

### Data and quality control

We mapped fine-grained genetic structure in the Netherlands using a population-based Dutch ALS case-control dataset (n=1,626; subset of stratum sNL3 from a genome-wide association study (GWAS) for amyotrophic lateral sclerosis^17^) and a European reference dataset subsampled from a GWAS for multiple sclerosis^19^ (MS; n = 4,514; EGA accession ID EGAD00000000120). 1,422 Dutch individuals had associated residential data (hometown at time of sampling) which were used for geographical analyses. For estimating GWAS confounding, we separately analysed the Netherlands on its own using a larger ALS case/control dataset (n = 4,753; strata sNL1, sNL3 and sNL4 from reference 17) and the complete multi-population GWAS dataset^17^ (n = 36,052) from which this Dutch subset was derived. Data handling for estimating confounding is further described under “Estimating GWAS confounding” below. For population structure analyses, we applied quality control (QC) using PLINK v1.9^31^; briefly we removed samples with high missingness (>10%), high heterozygosity (>3 median absolute deviations from median) and single-marker PCA outliers (>5 standard deviations from mean for PCs 1-20). We also filtered out A/T and G/C SNPs and SNPs with minor allele frequency <0.05, high missingness (>2%) or in Hardy Weinberg disequilibrium (p<1×10^−6^). Before running Chromopainter/fineSTRUCTURE we retained only one individual from any pair or group that exhibited greater than 7.5% genomic relatedness (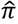) and removed SNPs with any missing genotypes as the algorithm does not tolerate missingness or relatedness well. For European reference data we also removed individuals suggested by the QC of the source study^19^ and we extracted individuals only of European descent. As this European dataset included MS patients, we filtered out SNPs in a 15 Mb region surrounding the strongly associated HLA locus (GRCh37 position chr6:22,915,594–37,945,593) to avoid bias generated from this association, following previous works. The final Dutch and European reference datasets contained 374,629 SNPs and 363,396 SNPs respectively at zero missingness. The merge of these datasets contained 147,097 SNPs at zero missingness. Data were phased per chromosome with the 1000 Genomes Project phase 3 reference panel^32^ using SHAPEIT v2^33^ (for ChromoPainter/fineSTRUCTURE) and Beagle v4.1 (for IBD estimation). Both programmes were run with default settings; allele concordance was checked prior to phasing (SHAPEIT: --check; Beagle: conform-gt utility).

### fineSTRUCTURE analysis

We used ChromoPainter/fineSTRUCTURE^18^ to detect fine-grained population structure using default settings. In brief, each individual was painted using all other individuals (-a 0 0), first estimating N_e_ and μ (switch rate and mutation rate) with 10 expectation-maximization iterations, then the model was finally run using these parameter estimates. The fineSTRUCTURE Markov chain Monte Carlo (MCMC) model was then run on the resulting coancestry matrix with two chains for 3,000,000 burnin and 1,000,000 sampling iterations, sampling every 10,000 iterations. We extracted the state with the maximum posterior probability and performed an additional 10,000 burnin iterations before inferring the final trees using both the climbtree and maximum concordance methods. For all subsequent analyses the maximum concordance tree was used.

### Cluster robustness

To assess the robustness of clustering in the Dutch data we calculated TVD^4^ and F_ST_. TVD is a distance metric for assessing the distinctness of pairs of clusters, calculated from the ChromoPainter chunklength matrix. TVD is calculated as the sum of the absolute differences between copying vectors for all pairs of clusters, where the copying vector for a given cluster *A* is a vector of the average lengths of DNA donated to individuals in *A* by all clusters. Intuitively, the TVD of two clusters reflects distance between those clusters in terms of haplotype sharing amongst all clusters, and is a meaningful method for assessing the effectiveness of fineSTRUCTURE clustering. To assess whether the observed clustering performed better than chance we permuted individuals between cluster pairs (maintaining cluster size) and calculated the number of permutations that exceeded our original TVD score for that pairing of clusters. We used 1,000 permutations where possible, and otherwise used the maximum number of unique permutations. P-values were calculated from the number of permutations greater than or equal to the observed TVD divided by the total permutations; all p-values were less than 0.001, indicating robust clustering. Finally we generated a TVD tree for k=16 by merging pairs of clusters with the lowest TVD successively using methods described previously^8^ (Supplementary figure 5). The tree was built in k-1 steps, with TVD recalculated at each step from the remaining populations. Branch lengths were scaled proportional to the TVD value of the corresponding pair of populations using adapted code from the original paper. To assess cluster differentiation independently of the ChromoPainter model, F_ST_ was calculated between Dutch clusters using PLINK 1.9. For comparison, we calculated F_ST_ between European countries present in reference 19.

### Ancestry profiles

We assessed the ancestral profile of Dutch samples in terms of a European reference made up of 4,514 European individuals^19^ from Belgium, Denmark, Finland, France, Germany, Italy, Norway, Poland, Spain and Sweden. European samples were first assigned to homogeneous genetic clusters using fineSTRUCTURE as in previous work^6^ to reduce noise in painting profiles. We then modelled each Dutch individual’s genome as a linear mixture of the European donor groups using ChromoPainter, and applied ancestry profile estimation as described previously^4^ and implemented in GLOBETROTTER^12^ (num.mixing.iterations: 0). This method estimates the proportion of DNA which is most closely shared with each individual from each donor group calculated from a normalised ChromoPainter chunklength output matrix, and then implements a multiple linear regression of the form

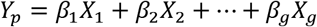

to correct for noise caused by similarities between donor populations. Here, *Y*_*p*_ is a vector of the proportion of DNA that individual *p* copies from each donor group, and *X*_*g*_ is the vector describing the average proportion of DNA that individuals in donor group *g* copy from other donor groups including their own. The coefficients of this equation *β*_1_… *β*_*g*_ are thus interpreted as the “cleaned” proportions of the genome that target individual *p* copies from each donor group, hence the ancestral contribution of each donor group to that individual. The equation is solved using a non-negative-least squares function such that *β*_*g*_ ≥ 0 and the sum of proportions across groups equals 1. We discarded European groups that contributed less than 5% total to any individual, and refit to eliminate noise. We then aggregated sharing proportions across donor groups (genetically homogenous clusters) from the same country to estimate total sharing between an individual and a given country to investigate the regional distribution of sharing profiles. Autocorrelation of ancestry profiles was assessed by Moran’s I and Mantel’s test (10,000 permutations) in R version 3.2.3. Geographical directions of ancestry gradients were determined by rotating the plane of latitude-longitude between 0° and 360° in 1° steps and finding the axis *Y* that maximised the coefficient of determination for the linear regression *Y*~*A*_*c*_, where *A*_*c*_ is the aggregated ancestry proportion for country *c*.

### Identity-by-descent analyses

IBD segments were called in phased data using RefinedIBD^20^ (default settings) to generate pairwise matrices of total length of IBD shared between individuals for bins of different segment lengths. To identify population structure captured by IBD sharing patterns we performed PCA on these matrices using the prcomp function in R version 3.2.3^34^ and clustered the IBD matrices using a Gaussian mixture model implemented in the R package mclust^35^. We note that while previous work^21^ has shown that IBD matrices underperform the linked ChromoPainter matrix in identifying population structure, they are arguably more interpretable for visualising temporal change as they can be subdivided into cM bins corresponding to different time periods, a feature leveraged by emerging work on local population structure^23^. Patterns in IBD sharing that identify population subgroups in older (shorter) cM bins which are preserved in more recent (longer) bins are interpreted as persistent population structure that has been influenced by mating patterns in old and recent generations. Structure which emerges in a specific cM bin and is lost is likely to reflect transient changes in panmixia that have not necessarily persisted. We approximated the age of segments in a given cM bin using equation s19 from reference 23, under the assumption that the population is sufficiently large.

### Inferring admixture events

To infer and date admixture events from European sources we ran GLOBETROTTER^12^ with the Netherlands dataset as a whole and in individual cluster groups defined from the Dutch fineSTRUCTURE maximum concordance tree (Figure 1). To define European donor groups we used the European fineSTRUCTURE maximum concordance tree, as with previous work^6^ to ensure genetically homogenous donor populations. We used ChromoPainter v2 to paint Dutch and European individuals using European clusters as donor groups. This generated a copying matrix (chunklengths file) and 10 painting samples for each Dutch individual. GLOBETROTTER was run for 5 mixing iterations twice: once using the null.ind:1 setting to test for evidence of admixture accounting for unusual linkage disequilibrium (LD) patterns and once using null.ind:0 to finally infer dates and sources. We further ran 100 bootstraps for the admixture date and calculated the probability of no admixture as the proportion of nonsensical inferred dates (<1 or >400 generations). Confidence intervals were calculated from the bootstraps from the standard model (null.ind:0) using the empirical bootstrap method, and a generation time of 28 years.

### ADMIXTURE analysis

We performed ADMIXTURE analysis^36^ on the combined Dutch and European samples to explore single marker-based population structure in a set of 41,675 SNPs (LD-pruned using PLINK 1.9: r^2^ > 0.1; sliding window 50 SNPs advancing 10 SNPs at a time) SNPs. ADMIXTURE was run for k=1-10 populations, using 5 EM iterations at each k value. The k value with the lowest cross validation error was selected for further analysis. We analysed the distribution of proportions for each ADMIXTURE cluster across the Dutch dataset, and its relationship with geography.

### Estimating mean pairwise IBD sharing within and between groups

We compared IBD sharing within and between both clusters and provinces (Supplementary figure 2) using the mean number of segments within a given length range (eg 1-2cM) shared between individuals. To calculate this mean for a single group of size *N* with itself the denominator was (*N*^2^ − *N*)/2; when comparing two groups of sizes *N* and *M* the denominator was *NM*.

### Estimating recent changes in population sizes

We used IBDNe^24^ to estimate historical changes in N_e_. IBDNe leverages information from the length distribution of IBD segments to accurately estimate effective population size over recent generations, with a resolution limit of about 50 generations for SNP data. We followed the authors’ protocol and detected IBD segments using IBDseq version r1206^37^ with default settings and ran IBDNe on the resulting output with default settings, removing IBD segments shorter than 4cM (minibd=4, the recommended threshold for genotype data). We compared estimated N_e_ with recorded census size (https://opendata.cbs.nl/statline/#/CBS/nl/dataset/37296ned/table?ts=1520261958200) for approximately equivalent dates (starting at 1946 CE for generation 0 and assuming 1 generation is 28 years) and found that for generations 0 - 3 our N_e_ estimates were approximately ⅓ of the census population (Supplementary figure 6), which follows expectation if lifespan is 3× the generation time. The slope of the ratios for the three generations is near zero suggesting that our model tracks well with the census population; this is consistent with reported expectation^24^.

### Estimating effective migration surfaces

To model geographic barriers to geneflow in the Netherlands we ran EEMS^14^. This software provides a visualisation of hot and coldspots for geneflow across a habitat using a geocoded genetic dataset. To run EEMS, we generated an average pairwise genetic dissimilarity matrix from our genotype data using the bed2diffs utility provided with the software. We initially ran the EEMS model with 10 randomly initialised MCMC chains for a short run of 100,000 burn-in and 200,000 sampling iterations, thinning every 999 iterations, to find a suitable starting point. For these runs we placed the data in 800 demes and used default settings with the following adjustments to the proposal variances: qEffctProposalS2 =0.00008888888; qSeedsProposalS2 = 0.7; mEffctProposalS2 = 0.7. The resulting chain with the highest log-likelihood was then used as the starting point for a further ten chains for 1,000,000 burn-in iterations and 2,000,000 sampling iterations, thinning every 9,999 iterations. The model was run with the following adjustments to the proposal variances: qEffctProposalS2 = 0.00008888888; qSeedsProposalS2 = 0.7; mEffctProposalS2 = 0.7. We plotted the results of our analysis using the rEEMSplot package in R and modified the resulting vector graphics using Inkscape v0.91 to remove display artefacts caused by non-overlapping polygons. MCMC convergence was assessed by inspecting the log-posterior traces (Supplementary figure 7).

### Estimating GWAS confounding

To examine the contribution of observed fine-grained population structure to GWAS confounding, we estimated how well phenotype could be predicted by principal components of haplotype sharing matrices in a 2016 GWAS for ALS^17^, comparing our results to those obtained using standard single marker PCA. We separately analysed 1,060,224 zero-missingness Hapmap3 SNPs that passed QC in the original GWAS for Dutch data alone (1,971 cases, 2,782 controls) and for the complete multi-population GWAS (12,577 cases, 23,475 controls). Haplotypes for unrelated individuals (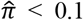) were phased using SHAPEIT v2^33^ and painted in terms of one another using ChromoPainter v2^18^ for the Dutch dataset (estimating N_e_ and μ using the weighted average of 10 EM iterations on chromosomes 1, 8, 15 and 20), and PBWT-paint (https://github.com/richarddurbin/pbwt/blob/master/pbwtPaint.c) for the considerably larger multi-population GWAS dataset. PBWT-paint is a fast approximate implementation of ChromoPainter suitable for large datasets. PCs of the resulting coancestry matrices were calculated using the fineSTRUCTURE R tools (https://www.paintmychromosomes.com). For comparison we also calculated PCs on independent markers from the SNP datasets using Plink v1.9, first removing long range LD regions^38^ (https://genome.sph.umich.edu/wiki/Regions_of_high_linkage_disequilibrium_(LD)) and pruning for LD (--indep-pairwise 500 50 0.8). Variance in ALS phenotype explained by ChromoPainter/PBWT-paint PCs and SNP PCs (Nagelkerke R^2^) was estimated using the glm() function and fmsb package^39^ in R version 3.2.3. To estimate confounding in GWAS inflation, we implemented a logistic regression model GWAS (--logistic) in PLINK v1.9 for each dataset using 20 ChromoPainter/PBWT-paint PCs or 20 SNP PCs as covariates and ran LD score regression^40^ on the resulting summary statistics using recommended settings. Structure evident in the PBWT-paint matrix was visualised and contrasted with corresponding SNP data in 2 dimensions using t-distributed stochastic neighbour embedding (t-SNE)^41^ implemented in the Rtsne package in R version 3.2.3 (5,000 iterations; perplexity 30; top 100 PCs provided as initial dimensions).

## Acknowledgements

This work has been supported by Science Foundation Ireland (17/CDA/4737), the Motor Neurone Disease Association of England, Wales and Northern Ireland (957-799) and the European Research Council (ERC) under the European Union’s Horizon 2020 research and innovation programme (grant agreement n° 772376 – EScORIAL). The collaboration project is co-funded by the PPP Allowance made available by Health~Holland, Top Sector Life Sciences & Health, to stimulate public-private partnerships.

## Author contributions

R.P.B. and R.L.McL. conceived the study. R.P.B, W.v.R., J.H.V and R.L.McL. contributed to study design. R.P.B. and R.L.McL. conducted the analyses. R.P.B. and R.L.McL. drafted the manuscript. W.v.R., L.H.v.d.B. and J.H.V. provided data and critical revision of the manuscript.

## Conflict of interest statement

All authors have nothing to declare.

## Supplementary material

**Supplementary table 1.** Mean pairwise F_ST_ (×10^−3^) for European groups from Sawcer *et al*.^19^

|  | **Finland** | **Sweden** | **Norway** | **Germany** | **Italy** | **Denmark** | **Belgium** | **Spain** | **Poland** | **France** |
| --- | --- | --- | --- | --- | --- | --- | --- | --- | --- | --- |
| **Finland** | 0 | 3.93 | 5.10 | 5.85 | 11.1 | 5.56 | 6.71 | 9.99 | 5.81 | 7.59 |
| **Sweden** | - | 0 | 0.362 | 0.702 | 4.87 | 0.362 | 1.07 | 3.76 | 1.84 | 1.81 |
| **Norway** | - | - | 0 | 0.899 | 4.96 | 0.407 | 1.07 | 3.73 | 2.56 | 1.76 |
| **Germany** | - | - | - | 0 | 2.77 | 0.399 | 0.289 | 2.11 | 1.26 | 0.678 |
| **Italy** | - | - | - | - | 0 | 4.08 | 2.29 | 1.21 | 5.42 | 1.55 |
| **Denmark** | - | - | - | - | - | 0 | 0.526 | 3.01 | 2.13 | 1.22 |
| **Belgium** | - | - | - | - | - | - | 0 | 1.53 | 2.43 | 0.367 |
| **Spain** | - | - | - | - | - | - | - | 0 | 4.56 | 0.660 |
| **Poland** | - | - | - | - | - | - | - | - | 0 | 2.84 |
| **France** | - | - | - | - | - | - | - | - | - | 0 |

**Supplementary figure 1.**
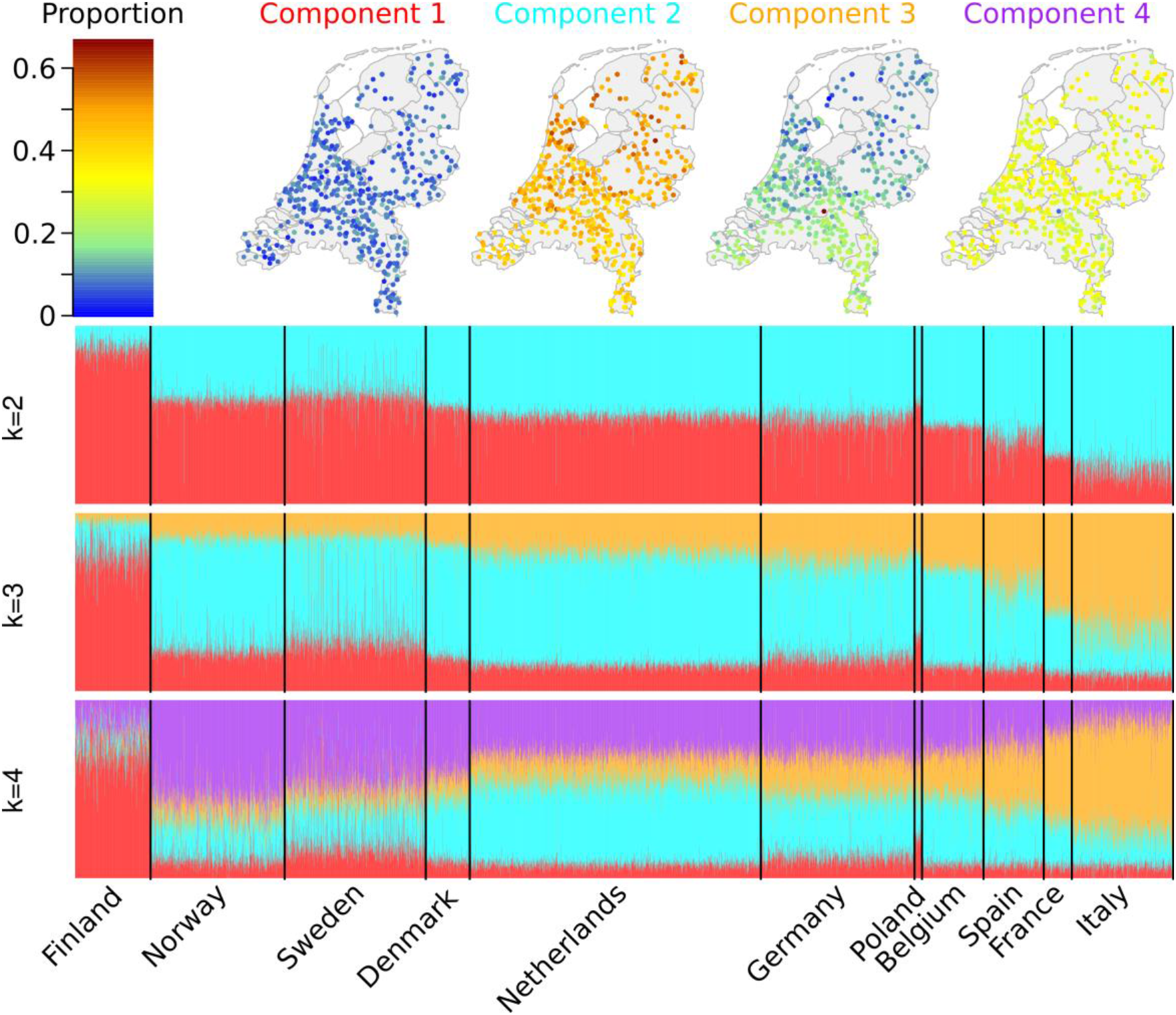
ADMIXTURE modelling for Dutch and European samples. Maps depict the regional breakdown of ADMIXTURE components for k=4 split. Dutch samples have a high value for admixture component 2, which is next highest in Germany and Belgium. Components 2 and 3 show opposing north-south gradients in the Netherlands, with component 2 highest in the north and component 3 highest in the south. Component 3 is best represented in southern European countries such as Italy.

**Supplementary figure 2.**
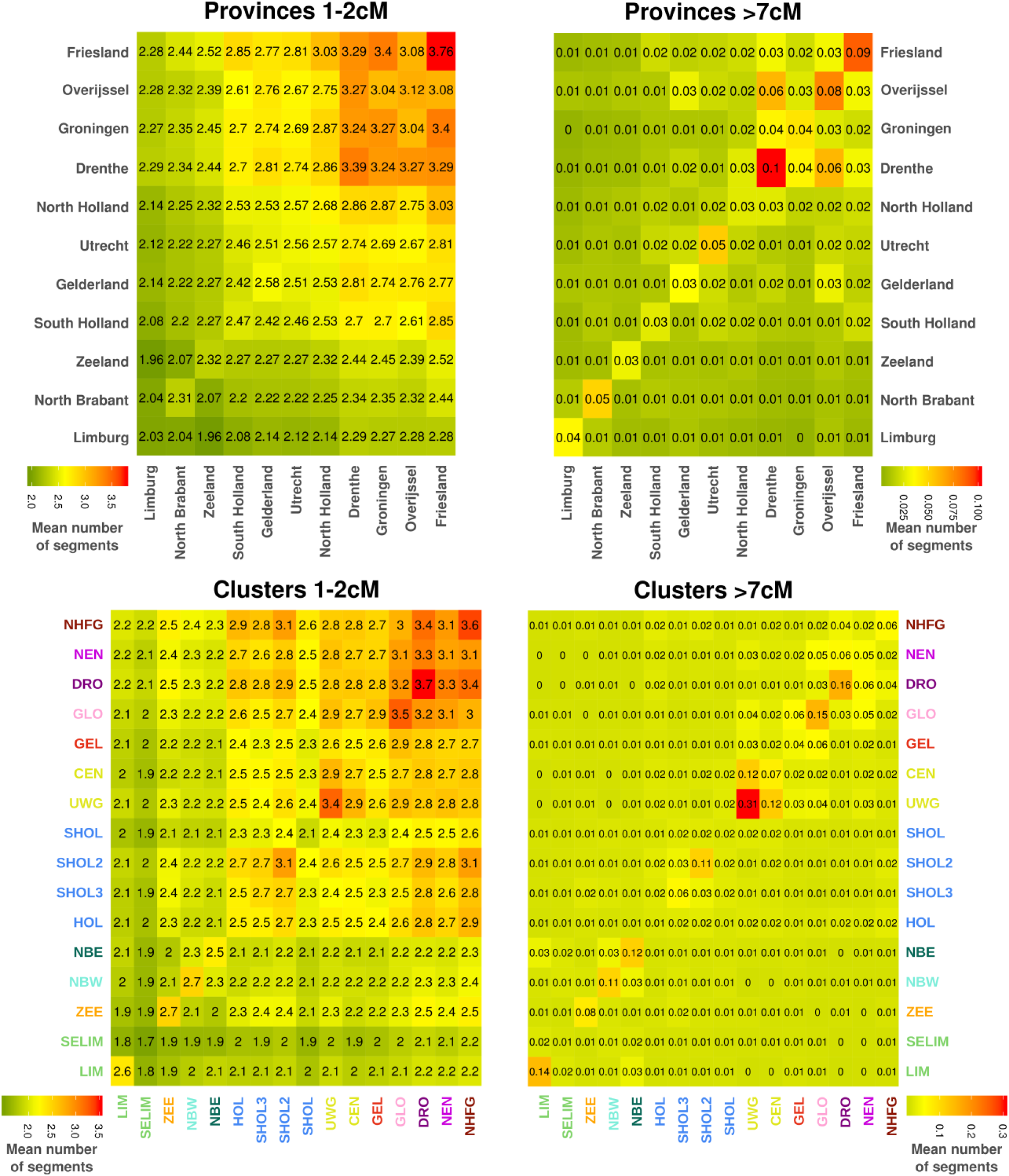
Old (left) and recent (right) IBD sharing per province (top) and per cluster (bottom). Average sharing of old (short) segments is enriched in northern provinces and clusters. Average sharing of recent (long) segments is higher on average within clusters than within provinces, indicating haplotypic clustering captures marginally more recent ancestry.

**Supplementary figure 3.**
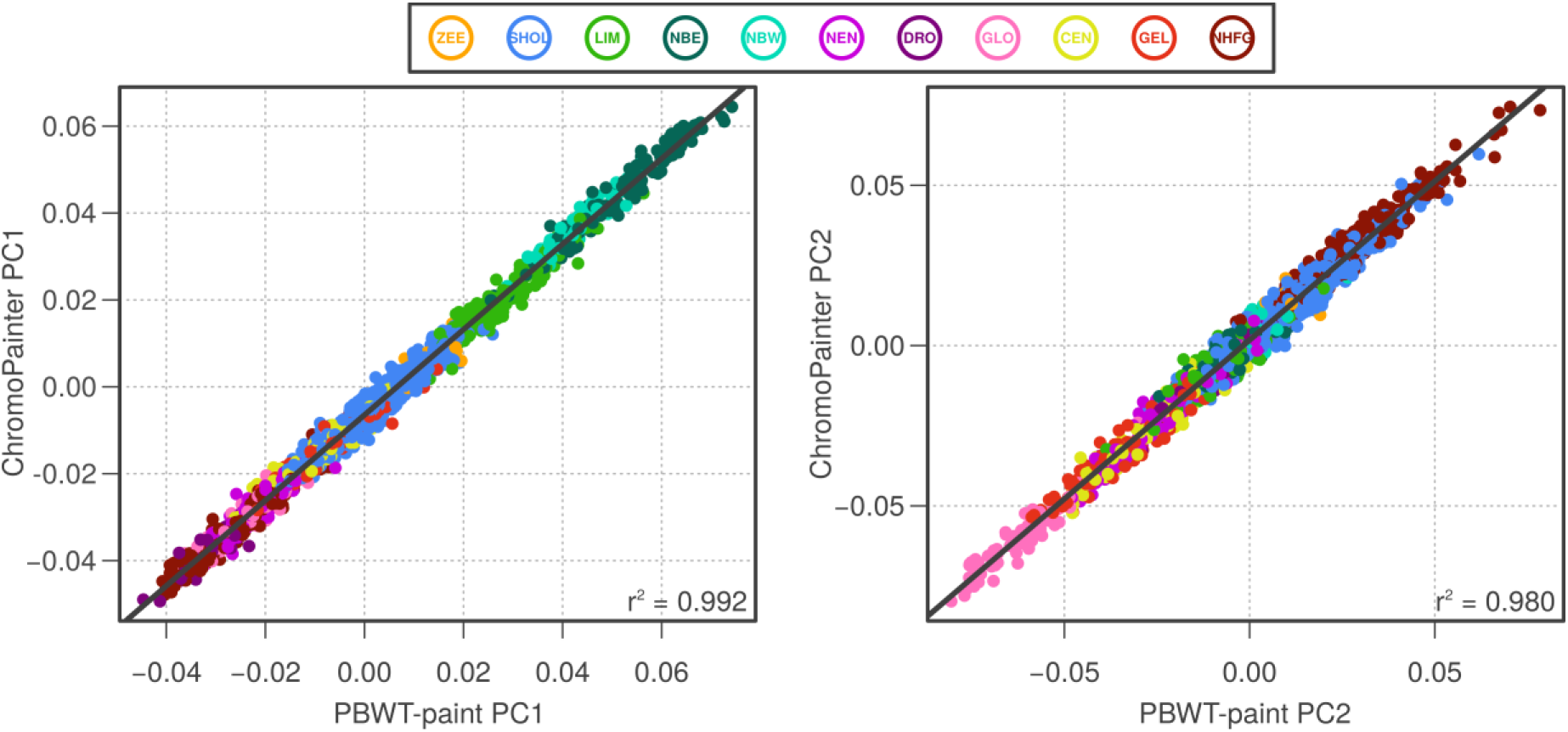
Benchmark of PBWT-paint vs ChromoPainter. Scatterplots comparing the first two principal components (PCs) of the coancestry matrices produced by ChromoPainter and PBWT-paint, showing strong correlation. Points are coloured by cluster groups defined in Figure 1. For all pairwise comparisons in the two coancestry matrices, Pearson’s ρ = 0.82 (0.82-0.821; p < 2×10^−16^).

**Supplementary figure 4.**
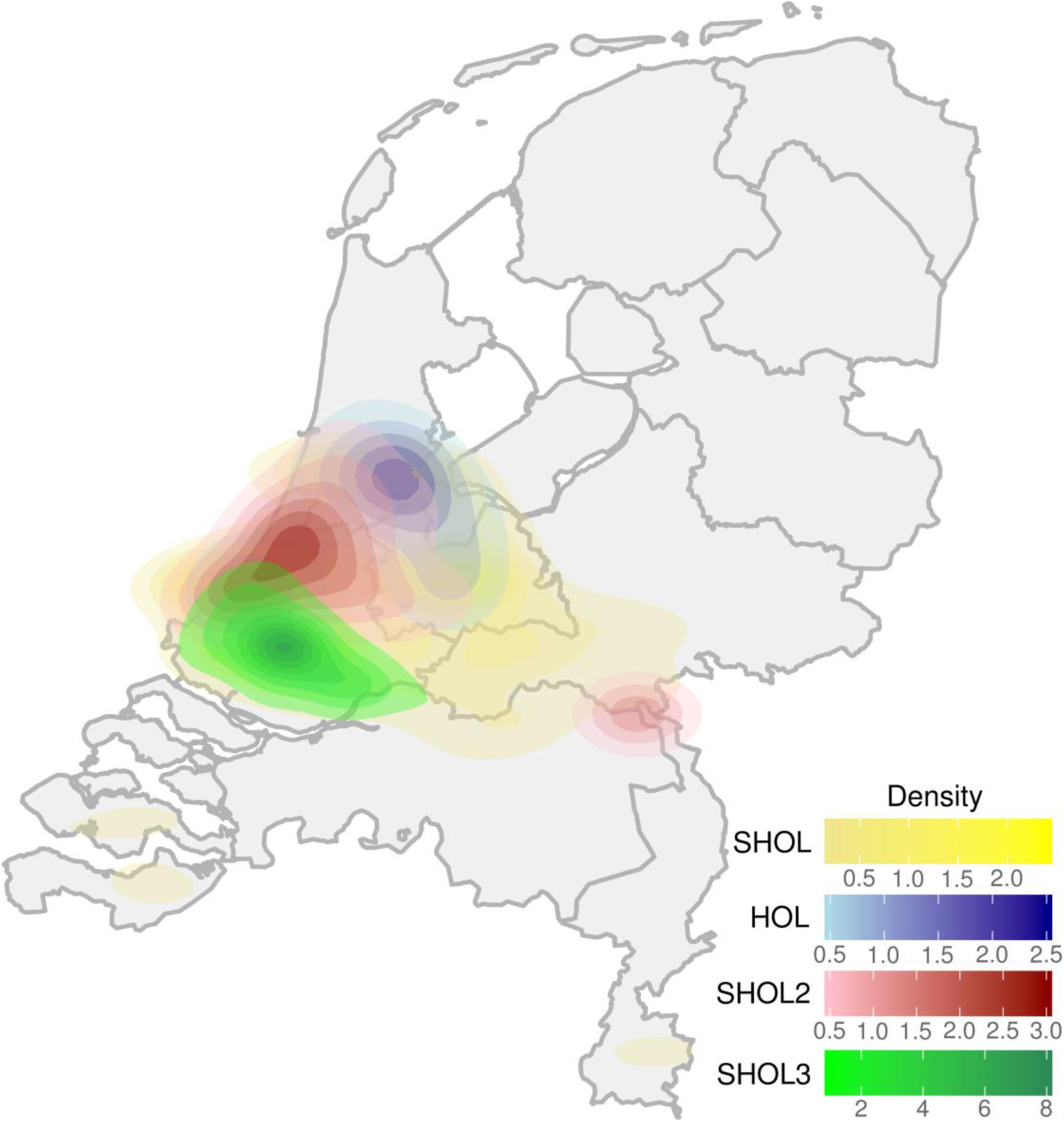
Geographic distribution of South Holland clusters from the SHOL cluster group. 2D kernel density estimates are shown for the geographic spread of samples from clusters SHOL (yellow), HOL (blue), SHOL2 (red), and SHOL3 (green) which form the SHOL cluster group in Figure 1. Kernel density estimates were calculated using the stat_density2d function in ggplot2 (R version 3.2.3) with default settings. >80% of samples are contained within plotted polygons for each cluster. Notably, although overlapping, three of the four clusters show quite distinct geographic ranges.

**Supplementary figure 5.**
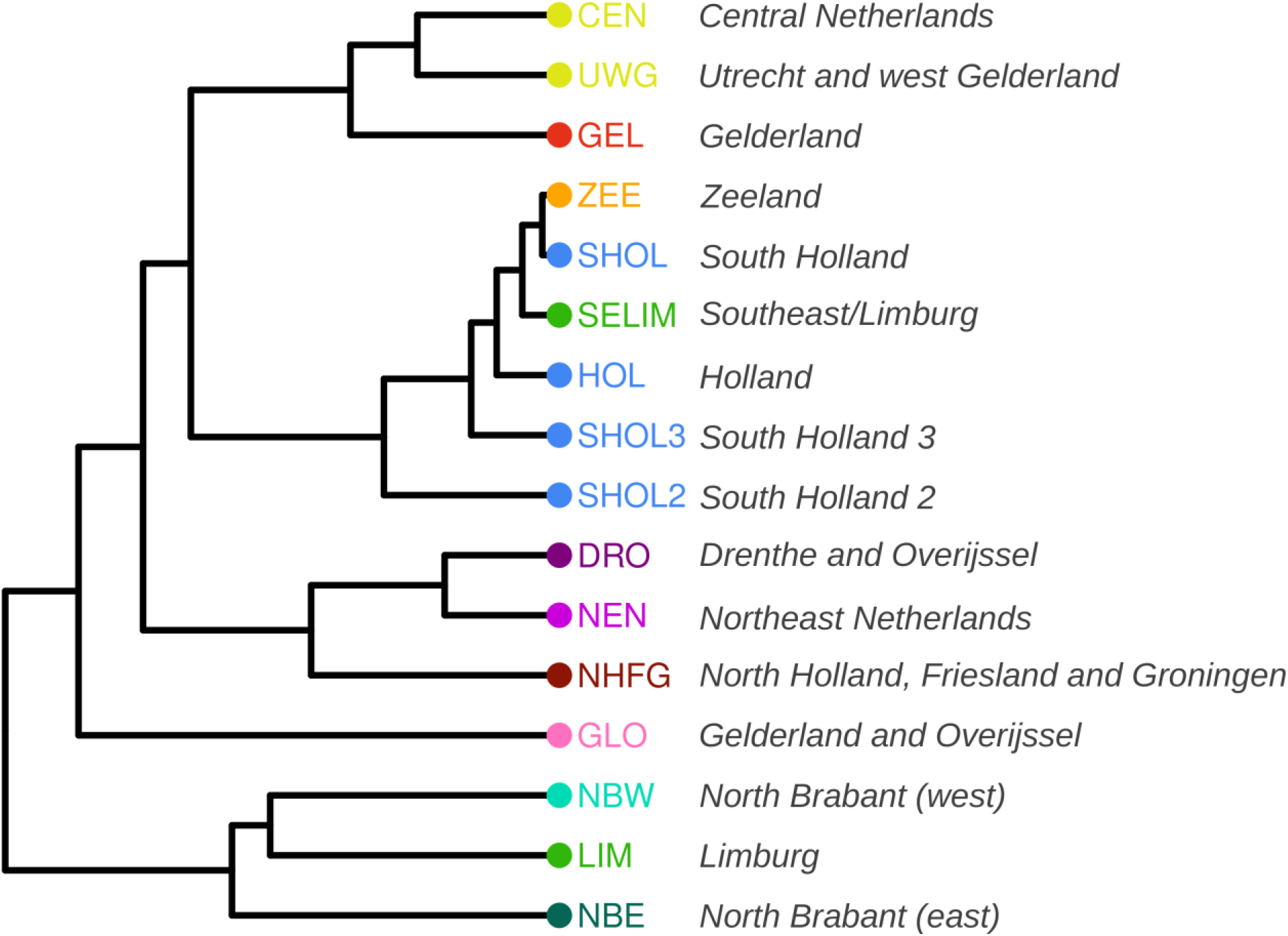
Total variation distance (TVD) tree for k=16 split in the Netherlands. Clusters are coloured and labelled according to scheme in Figure 1.

**Supplementary figure 6.**
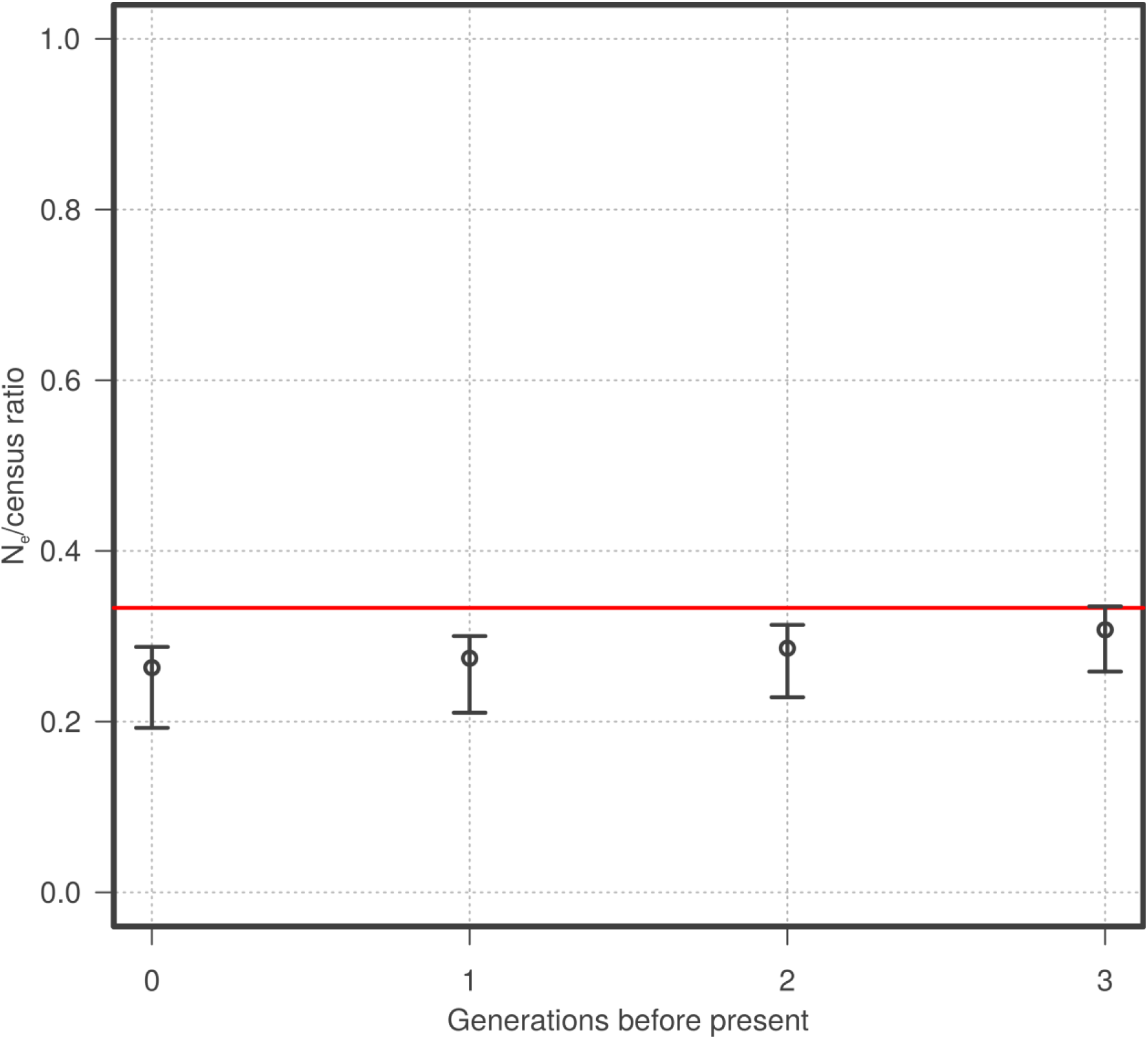
Ratio of estimated Ne/Census is stable over the past 3 generations. The red line at 0.33 corresponds to the expected ratio of N_e_ to census if lifespan is 3 times the generation time.

**Supplementary figure 7.**
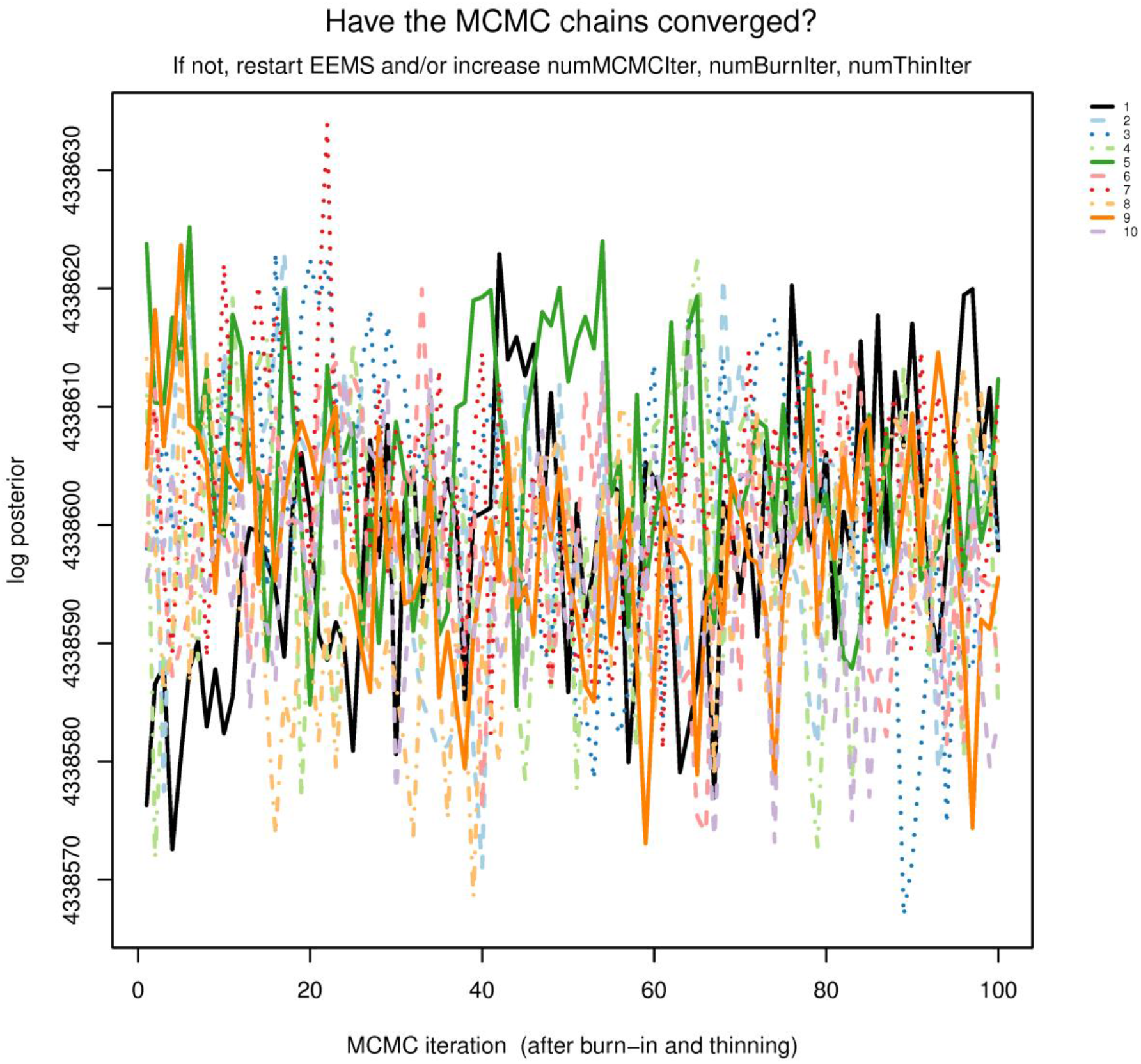
Convergence of MCMC chains for EEMS run in Netherlands. 10 independently seeded MCMC chains reach approximate convergence.

## Supplementary note 1

### Project MinE ALS GWAS Consortium authors

Wouter van Rheenen^1^, Aleksey Shatunov^2^, Russell L. McLaughlin^3^, Rick A.A. van der Spek^1^, Alfredo Iacoangeli^2,4^, Kevin P. Kenna^1^, Kristel R. van Eijk^1^, Nicola Ticozzi^5,6^, Boris Rogelj^7,8^, Katarina Vrabec^9^, Metka Ravnik-Glavač^9,10^, Blaž Koritnik^11^, Janez Zidar^11^, Lea Leonardis^11^, Leja Dolenc Grošelj^11^, Stéphanie Millecamps^12^, François Salachas^12,13,14^, Vincent Meininger^15,16^, Mamede de Carvalho^17,18^, Susana Pinto^17^, Marta Gromicho^17^, Ana Pronto-Laborinho^17^, Jesus S. Mora^19^, Ricardo Rojas-García^20,21^, Meraida Polak^22,23^, Siddharthan Chandran^24,25^, Shuna Colville^24^, Robert Swingler^24^, Karen E. Morrison^26^, Pamela J. Shaw^27^, John Hardy^28^, Richard W. Orrell^29^, Alan Pittman^28,30^, Katie Sidle^29^, Pietro Fratta^31^, Andrea Malaspina^32,33^, Simon Topp^2^, Susanne Petri^34^, Susanna Abdulla^35^, Carsten Drepper^36^, Michael Sendtner^36^, Thomas Meyer^37^, Roel A. Ophoff^38,39^, Kim A. Staats^39^, Martina Wiedau-Pazos^40^, Catherine Lomen-Hoerth^41^, Vivianna M. Van Deerlin^42^, John Q. Trojanowski^42^, Lauren Elman^43^, Leo McCluskey^43^, A. Nazli Basak^44^, Thomas Meitinger^45^, Peter Lichtner^45^, Milena Blagojevic-Radivojkov^45^, Christian R. Andres^46^, Gilbert Bensimon^47,48,49^, Bernhard Landwehrmeyer^50^, Alexis Brice^51,52,53,54,55^, Christine A.M. Payan^47,49^, Safaa Saker-Delye^56^, Alexandra Dürr^57^, Nicholas W. Wood^58^, Lukas Tittmann^59^, Wolfgang Lieb^59^, Andre Franke^60^, Marcella Rietschel^61^, Sven Cichon^62,63,64,65,66^, Markus M. Nöthen^62,63^, Philippe Amouyel^67^, Jean-François Dartigues^68^, Andre G. Uitterlinden^69,70^, Fernando Rivadeneira^69,70^, Karol Estrada^69^, Albert Hofman^70,71^, Charles Curtis^72,73^, Anneke J. van der Kooi^74^, Markus Weber^75^, Christopher E. Shaw^2^, Bradley N. Smith^2^, Daisy Sproviero^76^, Cristina Cereda^76^, Mauro Ceroni^77^, Luca Diamanti^77^, Roberto Del Bo^78^, Stefania Corti^78^, Giacomo P. Comi^78^, Sandra D’Alfonso^79^, Lucia Corrado^79^, Cinzia Bertolin^80^, Gianni Sorarù^80^, Letizia Mazzini^81^, Viviana Pensato^82^, Cinzia Gellera^82^, Cinzia Tiloca^5^, Antonia Ratti^5,6^, Andrea Calvo^83,84^, Cristina Moglia^83,84^, Maura Brunetti^83,84^, Rosa Capozzo^85^, Chiara Zecca^85^, Christian Lunetta^86^, Silvana Penco^87^, Nilo Riva^88^, Alessandro Padovani^89^, Massimiliano Filosto^90^, PARALS registry^91^, SLALOM group^91^, SLAP registry^91^, SLAGEN Consortium^91^, NNIPPS Study Group^91^, Ian Blair^92^, Garth A. Nicholson^92,93^, Dominic B. Rowe^92^, Roger Pamphlett^94^, Matthew C. Kiernan^95^, Julian Grosskreutz^96^, Otto W. Witte^96^, Robert Steinbach^96^, Tino Prell^96^, Beatrice Stubendorff^96^, Ingo Kurth^97,98^, Christian A. Hübner^97^, P. Nigel Leigh^99^, Federico Casale^83^, Adriano Chio^83,84^, Ettore Beghi^100^, Elisabetta Pupillo^100^, Rosanna Tortelli^101^, Giancarlo Logroscino^102,103^, John Powell^2^, Albert C. Ludolph^50^, Jochen H. Weishaupt^50^, Wim Robberecht^104,105,106^, Philip Van Damme^104,105,106^, Robert H. Brown^107^, Jonathan D. Glass^22,23^, John E. Landers^107^, Orla Hardiman^108,109^, Peter M. Andersen^50,110^, Philippe Corcia^46,111,112^, Patrick Vourc’h^46^, Vincenzo Silani^5,6^, Michael A. van Es^1^, R. Jeroen Pasterkamp^113^, Cathryn M. Lewis^72,114^, Gerome Breen^72,73^, Ammar Al-Chalabi^2^, Leonard H. van den Berg^1^, Jan H. Veldink^1^

1. Department of Neurology, UMC Utrecht Brain Center, University Medical Center Utrecht, Utrecht University, Utrecht, The Netherlands.

2. Maurice Wohl Clinical Neuroscience Institute, King’s College London, Department of Basic and Clinical Neuroscience, London, UK.

3. Complex Trait Genomics Laboratory, Smurfit Institute of Genetics, Trinity College Dublin, Dublin, Republic of Ireland.

4. Department of Biostatistics and Health Informatics, Institute of Psychiatry, Psychology and Neuroscience, King’s College London, London, UK.

5. Department of Neurology and Laboratory of Neuroscience, IRCCS Istituto Auxologico Italiano, Milano, Italy.

6. Department of Pathophysiology and Tranplantation, ‘Dino Ferrari’ Center, Università degli Studi di Milano, Milano, Italy.

7. Department of Biotechnology, Jožef Stefan Institute, Ljubljana, Slovenia.

8. Biomedical Research Institute BRIS, Ljubljana, Slovenia.

9. Department of Molecular Genetics, Institute of Pathology, Faculty of Medicine, University of Ljubljana, SI-1000 Ljubljana, Slovenia.

10. Institute of Biochemistry, Faculty of Medicine, University of Ljubljana, SI-1000 Ljubljana, Slovenia.

11. Ljubljana ALS Centre, Institute of Clinical Neurophysiology, University Medical Centre Ljubljana, SI-1000 Ljubljana, Slovenia.

12. Institut du Cerveau et de la Moelle épinière, Inserm U1127, CNRS UMR 7225, Sorbonne Universités, UPMC Univ Paris 06 UMRS1127, Paris, France.

13. Centre de Référence Maladies Rares SLA Ile de France, Département de Neurologie, Hôpital de la Pitié-Salpêtrière, Paris, France.

14. GRC-UPMC SLA et maladies du Motoneurone, France.

15. Ramsay Generale de Santé, Hopital Peupliers, Paris, France.

16. Réseau SLA Ile de France.

17. Instituto de Fisiologia, Instituto de Medicina Molecular,Faculdade de Medicina, Universidade de Lisboa, Lisbon, Portugal

18. Department of Neurosciences, Hospital de Santa Maria-CHLN, Lisbon, Portugal.

19. ALS Unit, Hospital San Rafael, Madrid, Spain

20. Neurology Department, Hospital de la Santa Creu i Sant Pau de Barcelona, Autonomous University of Barcelona, Barcelona, Spain.

21. Centro de Investigación en red en Enfermedades Raras (CIBERER), Spain.

22. Department Neurology, Emory University School of Medicine, Atlanta, GA, USA.

23. Emory ALS Center, Emory University School of Medicine, Atlanta, GA, USA.

24. Euan MacDonald Centre for Motor Neurone Disease Research, Edinburgh, UK.

25. Centre for Neuroregeneration and Medical Research Council Centre for Regenerative Medicine, University of Edinburgh, Edinburgh, UK.

26. School of Medicine, Dentistry and Biomedical Sciences, Queen’s University Belfast, UK.

27. Sheffield Institute for Translational Neuroscience (SITraN), University of Sheffield, Sheffield, UK.

28. Department of Molecular Neuroscience, Institute of Neurology, University College London, UK.

29. Department of Clinical Neuroscience, Institute of Neurology, University College London, UK.

30. Reta Lila Weston Institute, Institute of Neurology, University College London, UK.

31. Department of Neuromuscular Diseases, UCL Queen Square Institute of Neurology.

32. Centre for Neuroscience and Trauma, Blizard Institute, Queen Mary University of London, London, UK.

33. North-East London and Essex Regional Motor Neuron Disease Care Centre, London, UK.

34. Department of Neurology, Hannover Medical School, Hannover, Germany.

35. Department of Neurology, Otto-von-Guericke University Magdeburg, Magdeburg, Germany.

36. Institute of Clinical Neurobiology, University Hospital Wuerzburg, Germany.

37. Charité – Universitätsmedizin, Berlin, Germany.

38. Department of Human Genetics, David Geffen School of Medicine, University of California, Los Angeles, CA, USA.

39. Center for Neurobehavioral Genetics, Semel Institute for Neuroscience and Human Behavior, University of California, Los Angeles, CA, USA.

40. Department of Neurology, David Geffen School of Medicine, University of California, Los Angeles, CA, USA.

41. Department of Neurology, University of California, San Francisco, CA, USA.

42. Center for Neurodegenerative Disease Research, Perelman School of Medicine at the University of Pennsylvania, Philadelphia, PA, USA.

43. Department of Neurology, Perelman School of Medicine at the University of Pennsylvania, PA USA.

44. Koç University, School of Medicine, KUTTAM-NDAL, Istanbul Turkey.

45. Institute of Human Genetics, Helmholtz Zentrum München, Neuherberg, Germany.

46. Centre SLA, CHRU de Tours, Tours, France; UMR 1253, iBrain, Université de Tours, Inserm, Tours, France.

47. APHP, Département de Pharmacologie Clinique, Hôpital de la Pitié-Salpêtrière, France.

48. Université Pierre & Marie Curie, Pharmacologie, Paris VI, Paris, France.

49. BESPIM, CHU-Nîmes, Nîmes, France.

50. Department of Neurology, Ulm University, Ulm, Germany.

51. INSERM U 1127, Hôpital de la Pitié-Salpêtrière, 75013 Paris, France.

52. CNRS UMR 7225, Hôpital de la Pitié-Salpêtrière, 75013 Paris, France.

53. Sorbonne Universités, Université Pierre et Marie Curie Paris 06 UMRS 1127, Hôpital de la Pitié-Salpêtrière, 75013 Paris, France.

54. Institut du Cerveau et de la Moelle épinière, Hôpital de la Pitié-Salpêtrière, 75013 Paris, France.

55. APHP, Département de Génétique, Hôpital de la Pitié-Salpêtrière, 75013 Paris, France.

56. Genethon, CNRS UMR 8587 Evry, France.

57. Department of Medical Genetics, l’Institut du Cerveau et de la Moelle Épinière, Hoptial Salpêtrière, 75013 Paris, France.

58. Department of Neurogenetics, Institute of Neurology, University College London, UK.

59. PopGen Biobank and Institute of Epidemiology, Christian Albrechts-University Kiel, Kiel, Germany.

60. Institute of Clinical Molecular Biology, Kiel University, Kiel, Germany.

61. Department of Genetic Epidemiology in Psychiatry, Central Institute of Mental Health, Faculty of Medicine Mannheim, University of Heidelberg, Germany

62. Institute of Human Genetics, University of Bonn, Bonn, Germany.

63. Department of Genomics, Life and Brain Center, Bonn, Germany.

64. Division of Medical Genetics, University Hospital Basel, University of Basel, Basel, Switzerland.

65. Department of Biomedicine, University of Basel, Basel, Switzerland.

66. Institute of Neuroscience and Medicine INM-1, Research Center Juelich, Juelich, Germany.

67. University of Lille, Inserm, CHU Lille, Institut Pasteur de Lille, U1167 - RID-AGE - Risk Factor and molecular determinants of aging diseases, Labex Distalz, F-59000 Lille, France.

68. Bordeaux University, ISPED, Centre INSERM U1219-Epidemiologie-Biostatistique & CIC-1401, CHU de Bordeaux, Pole de Sante Publique, Bordeaux, France.

69. Department of Internal Medicine, Genetics Laboratory, Erasmus Medical Center Rotterdam, Rotterdam, The Netherlands.

70. Department of Epidemiology, Erasmus Medical Center Rotterdam, Rotterdam, The Netherlands.

71. Department of Epidemiology, Harvard T.H. Chan School of Public Health, Boston, MA, USA.

72. Social, Genetic & Developmental Psychiatry Centre, Institute of Psychiatry, Psychology & Neuroscience, King’s College London, London, UK.

73. NIHR Maudsley Biomedical Research Centre, Maudsley Hospital and Institute of Psychiatry, Psychology & Neuroscience, King’s College London, London, UK.

74. Amsterdam UMC, department of Neurology, University of Amsterdam, Neuroscience, Amsterdam

75. Neuromuscular Diseases Unit/ALS Clinic, Kantonsspital St. Gallen, 9007, St. Gallen, Switzerland.

76. Genomic and post-Genomic Center, IRCCS Mondino Foundation, Pavia, Italy.

77. General Neurology, IRCCS Mondino Foundation, Pavia, Italy

78. Neurologic Unit, IRCCS Foundation Ca’ Granda Ospedale Maggiore Policlinico, Milan, Italy.

79. Department of Health Sciences, Interdisciplinary Research Center of Autoimmune Diseases, UPO, Università del Piemonte Orientale, Novara, Italy.

80. Department of Neurosciences, University of Padova, Padova, Italy.

81. ALS Centre Department of Neurology Maggiore della Carità University Hospital, Novara.

82. Unit of Genetics of Neurodegenerative and Metabolic Diseases, Fondazione IRCCS Istituto Neurologico ‘Carlo Besta’, Milano, Italy.

83. “Rita Levi Montalcini” Department of Neuroscience, ALS Centre, University of Torino, Turin, Italy.

84. Azienda Ospedaliera Città della Salute e della Scienza, Torino, Italy.

85. Department of Clinical research in Neurology, University of Bari “A. Moro”, at Pia Fondazione “Card. G. Panico”, Tricase (LE), Italy.

86. NEMO Clinical Center, Serena Onlus Foundation, Niguarda Ca’ Granda Hostipal, Milan, Italy.

87. Medical Genetics Unit, Department of Laboratory Medicine, Niguarda Ca’ Granda Hospital, Milan, Italy.

88. Department of Neurology, Institute of Experimental Neurology (INSPE), Division of Neuroscience, San Raffaele Scientific Institute, Milan, Italy.

89. Neurology Unit, Department of Clinical and Experimental Sciences, University of Brescia, Italy.

90. Neurology Unit, Department of Neuroscience and Vision, Spedali Civili Hospital, Brescia, Italy.

91. A list of members and affiliations appears at the end of this Supplementary Note.

92. Department of Biomedical Sciences, Faculty of Medicine and Health Sciences, Macquarie University, Sydney, New South Wales, Australia.

93. University of Sydney, ANZAC Research Institute, Concord Hospital, Sydney, New South Wales, Australia.

94. Discipline of Pathology, Sydney Medical School, Brain and Mind Centre, The University of Sydney, New South Wales 2050, Australia.

95. Brain and Mind Centre, The University of Sydney, New South Wales 2050, Australia.

96. Hans-Berger Department of Neurology, Jena University Hospital, Jena, Germany.

97. Institute of Human Genetics, Jena University Hospital, Jena, Germany.

98. Institute of Human Genetics, Medical Faculty, RWTH Aachen University, Aachen, Germany

99. Department of Neurology, Brighton and Sussex Medical School Trafford Centre for Biomedical Research, University of Sussex, Falmer, East Sussex, UK.

100. Laboratory of Neurological Diseases, Department of Neuroscience, IRCCS Istituto di Ricerche Farmacologiche Mario Negri, Milano, Italy.

101. Institute of Neurology, University College of London (UCL), London, UK

102. Department of Basic Medical Sciences, Neuroscience and Sense Organs, University of Bari ‘Aldo Moro’, Bari, Italy.

103. Unit of Neurodegenerative Diseases, Department of Clinical Research in Neurology, University of Bari ‘Aldo Moro’, at Pia Fondazione Cardinale G. Panico, Tricase, Lecce, Italy.

104. KU Leuven - University of Leuven, Department of Neurosciences

105. VIB, Center for Brain & Disease Research, Laboratory of Neurobiology, Leuven, Belgium.

106. University Hospitals Leuven, Department of Neurology, Leuven, Belgium.

107. Department of Neurology, University of Massachusetts Medical School, Worcester, MA, USA.

108. Academic Unit of Neurology, Trinity College Dublin, Trinity Biomedical Sciences Institute, Dublin, Republic of Ireland.

109. Department of Neurology, Beaumont Hospital, Dublin, Republic of Ireland.

110. Department of Clinical Science, Neurosciences, Umeå University, Umeå, Sweden.

111. Federation des Centres SLA Tours and Limoges, LITORALS, Tours, France.

112. INSERM U1253, “iBrain”, Université François-Rabelais de Tours, Faculté de Médecine 10, Bd Tonnellé, 37032 Tours Cedex 1, France.

113. Department of Translational Neuroscience, UMC Utrecht Brain Center, University Medical Center Utrecht, Utrecht University, Utrecht, The Netherlands.

114. Department of Medical and Molecular Genetics, King’s College London, London, UK.

### Italian Consortium for the Genetics of ALS (SLAGEN) members

Daniela Calini, Isabella Fogh, Antonia Ratti, Vincenzo Silani, Nicola Ticozzi, Cinzia Tiloca, Barbara Castellotti, Cinzia Gellera, Viviana Pensato, Franco Taroni, Cristina Cereda, Mauro Ceroni, Stella Gagliardi, Giacomo Comi, Stefania Corti, Roberto Del Bo, Lucia Corrado, Sandra D’Alfonso, Letizia Mazzini, Elena Pegoraro, Giorgia Querin, Massimiliano Filosto and Gianni Sorarù

### Registro Lombardo Sclerosi Laterale Amyotrofica (SLALOM) group members

Francesca Gerardi, Fabrizio Rinaldi, Maria Sofia Cotelli, Luca Chiveri, Maria Cristina Guaita, Patrizia Perrone, Giancarlo Comi, Carlo Ferrarese, Lucio Tremolizzo, Marialuisa Delodovici, Massimiliano Filosto and Giorgio Bono

### Piemonte and Valle d’Aosta Registry for Amyotrophic Lateral Sclerosis (PARALS) group members

Stefania Cammarosano, Antonio Canosa, Dario Cocito, Leonardo Lopiano, Luca Durelli, Bruno Ferrero, Antonio Bertolotto, Alessandro Mauro, Luca Pradotto, Roberto Cantello, Enrica Bersano, Dario Giobbe, Maurizio Gionco, Daniela Leotta, Lucia Appendino, Roberto Cavallo, Enrico Odddenino, Claudio Geda, Fabio Poglio, Paola Santimaria, Umberto Massazza, Antonio Villani, Roberto Conti, Fabrizio Pisano, Mario Palermo, Franco Vergnano, Paolo Provera, Maria Teresa Penza, Marco Aguggia, Nicoletta Di Vito, Piero Meineri, Ilaria Pastore, Paolo Ghiglione, Danilo Seliak, Nicola Launaro, Giovanni Astegiano and Bottacchi Edo

### Sclerosi Laterale Amyotrofica-Puglia (SLAP) registry members

Isabella Laura Simone, Stefano Zoccolella, Michele Zarrelli and Franco Apollo

### Neuroprotection and Natural History in Parkinson Plus Syndromes (NNIPPS) Study group members

William Camu, Jean Sebastien Hulot, Francois Viallet, Philippe Couratier, David Maltete, Christine Tranchant, Marie Vidailhet.

## References

1. Abdellaoui, A. et al. Population structure, migration, and diversifying selection in the Netherlands. Eur J Hum Genet 21, 1277–1285 (2013).

2. Genome of the Netherlands Consortium. Whole-genome sequence variation, population structure and demographic history of the Dutch population. Nat. Genet. 46, 818–825 (2014).

3. Lawson, D. J. et al. Is population structure in the genetic biobank era irrelevant, a challenge, or an opportunity? Hum. Genet. (2019) doi:10.1007/s00439-019-02014-8.

4. Leslie, S. et al. The fine-scale genetic structure of the British population. Nature 519, 309–314 (2015).

5. Gilbert, E. et al. The Irish DNA Atlas: Revealing Fine-Scale Population Structure and History within Ireland. Sci Rep 7, 17199 (2017).

6. Byrne, R. P. et al. Insular Celtic population structure and genomic footprints of migration. PLoS Genet. 14, e1007152 (2018).

7. Gilbert, E. et al. The genetic landscape of Scotland and the Isles. Proc Natl Acad Sci U A 116, 19064–19070 (2019).

8. Kerminen, S. et al. Fine-Scale Genetic Structure in Finland. G3 7, 3459–3468 (2017).

9. Takeuchi, F. et al. The fine-scale genetic structure and evolution of the Japanese population. PLoS One 12, e0185487 (2017).

10. Raveane, A. et al. Population structure of modern-day Italians reveals patterns of ancient and archaic ancestries in Southern Europe. Sci Adv 5, eaaw3492 (2019).

11. Bycroft, C. et al. Patterns of genetic differentiation and the footprints of historical migrations in the Iberian Peninsula. Nat Commun 10, 551 (2019).

12. Hellenthal, G. et al. A genetic atlas of human admixture history. Science 343, 747–751 (2014).

13. Novembre, J. & Peter, B. M. Recent advances in the study of fine-scale population structure in humans. Curr Opin Genet Dev 41, 98–105 (2016).

14. Petkova, D., Novembre, J. & Stephens, M. Visualizing spatial population structure with estimated effective migration surfaces. Nat Genet 48, 94–100 (2016).

15. Buniello, A. et al. The NHGRI-EBI GWAS Catalog of published genome-wide association studies, targeted arrays and summary statistics 2019. Nucleic Acids Res. 47, D1005–D1012 (2019).

16. Mathieson, I. & McVean, G. Differential confounding of rare and common variants in spatially structured populations. Nat. Genet. 44, 243–246 (2012).

17. van Rheenen, W. et al. Genome-wide association analyses identify new risk variants and the genetic architecture of amyotrophic lateral sclerosis. Nat. Genet. 48, 1043–1048 (2016).

18. Lawson, D. J., Hellenthal, G., Myers, S. & Falush, D. Inference of population structure using dense haplotype data. PLoS Genet. 8, e1002453 (2012).

19. Sawcer, S. et al. Genetic risk and a primary role for cell-mediated immune mechanisms in multiple sclerosis. Nature 476, 214–219 (2011).

20. Browning, B. L. & Browning, S. R. Improving the accuracy and efficiency of identity-by-descent detection in population data. Genetics 194, 459–471 (2013).

21. Lawson, D. J. & Falush, D. Population identification using genetic data. Annu Rev Genomics Hum Genet 13, 337–361 (2012).

22. Palamara, P. F. Population genetics of identity by descent. (2014).

23. Al-Asadi, H., Petkova, D., Stephens, M. & Novembre, J. Estimating recent migration and population-size surfaces. PLoS Genet 15, e1007908 (2019).

24. Browning, S. R. & Browning, B. L. Accurate Non-parametric Estimation of Recent Effective Population Size from Segments of Identity by Descent. Am J Hum Genet 97, 404–418 (2015).

25. Herlihy, D. The Black Death and the Transformation of the West. (Harvard University Press, 1997).

26. Durbin, R. Efficient haplotype matching and storage using the positional Burrows-Wheeler transform (PBWT). Bioinforma. Oxf. Engl. 30, 1266–1272 (2014).

27. Athanasiadis, G. et al. Nationwide Genomic Study in Denmark Reveals Remarkable Population Homogeneity. Genetics 204, 711–722 (2016).

28. Nalls, M. A. et al. Measures of autozygosity in decline: globalization, urbanization, and its implications for medical genetics. PLoS Genet. 5, e1000415 (2009).

29. Roosen, J. & Curtis, D. R. The ‘light touch’of the Black Death in the Southern Netherlands: an urban trick? Econ Hist Rev 72, 32–56 (2019).

30. Haworth, S. et al. Apparent latent structure within the UK Biobank sample has implications for epidemiological analysis. Nat. Commun. 10, 333 (2019).

31. Chang, C. C. et al. Second-generation PLINK: rising to the challenge of larger and richer datasets. Gigascience 4, 7 (2015).

32. 1000 Genomes Project Consortium et al. A global reference for human genetic variation. Nature 526, 68–74 (2015).

33. Delaneau, O., Marchini, J. & Zagury, J.-F. cois. A linear complexity phasing method for thousands of genomes. Nat Methods 9, 179–181 (2011).

34. CoreTeam, R. R: A Language and Environment for Statistical Computing. Vienna, Austria: R Foundation for Statistical Computing; 2015. (2015).

35. Scrucca, L., Fop, M., Murphy, T. B. & Raftery, A. E. mclust 5: Clustering, Classification and Density Estimation Using Gaussian Finite Mixture Models. R J 8, 289–317 (2016).

36. Alexander, D. H., Novembre, J. & Lange, K. Fast model-based estimation of ancestry in unrelated individuals. Genome Res. 19, 1655–1664 (2009).

37. Browning, B. L. & Browning, S. R. Detecting identity by descent and estimating genotype error rates in sequence data. Am J Hum Genet 93, 840–851 (2013).

38. Price, A. L. et al. Long-range LD can confound genome scans in admixed populations. Am J Hum Genet 83, 132–5; author reply 135–9 (2008).

39. Nakazawa, M. fmsb: Functions for medical statistics book with some demographic data, 2014. R Package (2018).

40. Bulik-Sullivan, B. K. et al. LD Score regression distinguishes confounding from polygenicity in genome-wide association studies. Nat. Genet. 47, 291–295 (2015).

41. van der Maaten, L. & Hinton, G. Visualizing data using t-SNE. J. Mach. Learn. Res. 9, 2579–2605 (2008).

